# A non-canonical EZH2 function sensitizes solid tumors to genotoxic stress

**DOI:** 10.1101/2020.09.11.291534

**Authors:** Yiji Liao, Chen-Hao Chen, Neel Shah, Tengfei Xiao, Avery Feit, Mei Yang, Changmeng Cai, Shuai Gao, Pengya Xue, Zhijie Liu, Han Xu, Jihoon Lee, Wei Li, Shenglin Mei, Roodolph S. Pierre, Shaokun Shu, Teng Fei, Melissa Duarte, Jin Zhao, James E. Bradner, Kornelia Polyak, Philip W. Kantoff, Henry Long, Steven P. Balk, X. Shirley Liu, Myles Brown, Kexin Xu

**Author notes:** Correspondence: Xiaole Shirley Liu, Dana-Farber Cancer Institute, 450 Brookline Avenue, Boston, MA 02215.; (X.S.L.); Myles A. Brown, Dana-Farber Cancer Institute, 450 Brookline Avenue, Boston, MA 02215.; Kexin Xu, University of Texas Health Science Center at San Antonio, San Antonio, TX 78229. These authors contributed equally to this work (Yiji Liao, Chen-Hao Chen).

## Abstract

Drugs that block the activity of the methyltransferase EZH2 are in clinical development for the treatment of non-Hodgkin lymphomas harboring gain-of-function EZH2 mutations that enhance its polycomb repressive function. In contrast, in castration-resistant prostate cancer (CRPC) we have previously reported that EZH2 plays a non-canonical role as a transcriptional activator. In this setting, we now show that EZH2 inhibitors can also block the non-canonical activity of EZH2 and inhibit the growth of CRPC cells. Gene expression and epigenomic profiling of cells treated with EZH2 inhibitors demonstrated that rather than de-repressing tumor suppressor genes silenced by PRC2, EZH2 inhibitors downregulate a set of DNA repair genes that are directly regulated by EZH2. In addition, genome-wide CRISPR/Cas9-mediated loss-of-function screens in the presence of EZH2 inhibitors identified these DNA repair genes to underlie the growth-inhibitory function of these compounds. Interrogation of public data from diverse solid tumor types expressing wild-type EZH2 showed that expression of DNA damage repair genes is significantly correlated with cellular sensitivity to EZH2 inhibitors. Consistent with these findings, treatment of CRPC cells with EZH2 inhibitors dramatically enhanced their sensitivity to genotoxic stress. These studies reveal a previously unappreciated mechanism of action of EZH2 inhibitors and provide a mechanistic basis for potential new combination cancer therapies.

## Introduction

The methyltransferase EZH2 has shown encouraging therapeutic potential in cancer (Han Li and Chen, 2015). Originally identified as the catalytic subunit of the polycomb repressive complex 2 (PRC2), EZH2 methylates histone H3 at lysine 27 (H3K27) and leads to gene silencing (Cao et al., 2002). EZH2 is frequently upregulated in a broad spectrum of aggressive solid tumors and its overabundance is significantly associated with poor prognosis (Kim and Roberts, 2016). Gain-of-function mutations at residues Y641, A677 or A687 within the catalytic domain of EZH2 have been identified in diffuse large B-cell lymphoma (DLBCL) and follicular lymphoma (FL) (McCabe et al., 2012a; Morin et al., 2010). In view of these oncogenic features of EZH2, several selective inhibitors that block its enzymatic activity were developed (Knutson et al., 2012; McCabe et al., 2012b; Qi et al., 2012). These compounds specifically inhibit EZH2-mediated methyl transfer reactions by competing with the methyl donor S-adenosylmethionine (SAM) for the binding pocket inside the catalytic domain. These prototype drugs abrogated the growth of non-Hodgkin lymphoma (NHL) cells that harbor EZH2 driver mutations, decreased global tri-methylation of H3K27 (H3K27me3) and re-activated genes that are repressed by PRC2 complex. However, it remains unclear whether the efficacy of EZH2 inhibitors will be limited to NHL harboring gain-of-function mutations or will be active as well on a range of solid tumors that rarely contain somatic mutations of EZH2.

Genotoxic stress, such as that induced by radiation or chemotherapy, predisposes cells to DNA damages and elicits diverse biological responses, including DNA repair, cell cycle arrest and apoptosis (Swift and Golsteyn, 2014). Deregulation of components critical for an appropriate DNA damage response (DDR) leads to genome instability, a hallmark of most cancers. Therefore, drugs that induce DNA damage or inhibit DDR, such as cisplatin and PARP inhibitors, are effective anticancer agents across a wide array of tumor types. Growing evidence suggests that EZH2 plays a pivotal role in determining how cancer cells respond to DNA damage. In one study, knockdown of EZH2 predominantly induced apoptosis in both p53-proficient and -deficient cancer cells. This was dependent on H3K27me3-mediated epigenetic silencing of FBXO32, which is required for p21 protein degradation (Wu et al., 2011). In another report, depletion of EZH2 rapidly prompted a senescence-related DDR via activation of ATM-p53-p21 pathway. Interestingly, no changes in H3K27me3 pattern or overall level were observed during the process (Ito et al., 2018). In addition, EZH2 was found to be essential for cell proliferation following escape from senescence, but again changes in H3K27me3 was not involved in this phenotypic adaptation (Le Duff et al., 2018). Provocatively, it was recently reported that EZH2 directly binds to the internal ribosome entry site (IRES) on the mRNAs of both wild-type and mutated p53, leading to enhanced translation of p53 protein, especially mutant one, and subsequently promoted cancer growth and metastasis. However, this EZH2-RNA interaction seems to be independent of EZH2’s methyltransferase activity (Zhao et al., 2019). These studies present a diverse and complex picture of EZH2 functions in regulation of DDR. Additionally, the role of H3K27me3 in the EZH2-mediated DDR is unclear, raising the question of how exactly EZH2 triggers the specific biological responses to DNA damage. Conversely, it is largely unknown whether components involved in the DDR, in return, influence the oncogenic functions of EZH2. Unraveling answers to these questions will have a profound impact on the development of EZH2 inhibitors and combination cancer therapies.

In this study, we comprehensively evaluated and confirmed the therapeutic potentials of EZH2 inhibition in solid tumors using prostate cancer cell models. Interestingly, we found a non-canonical mechanism of action of EZH2 inhibitors in cancers without EZH2 somatic mutations. Rather than derepressing tumor suppressor genes silenced by PRC2-catalyzed H3K27m3, these compounds downregulate a set of DNA repair genes that are directly activated by EZH2. The same set of DNA repair genes is identified in CRISPR-Cas9 knockout screen to underlie the growth inhibitory function of EZH2 inhibitors, and well predicts sensitivities to EZH2 inhibitors across hundreds of cancer cell lines that express wild-type EZH2. More importantly, EZH2i potentiated the activity of DNA damaging agents and synergistically blocked the growth of advanced prostate cancer cells. Therefore, our work elucidates a novel mechanism of action of EZH2 inhibitors in solid tumors and suggests potential novel combination therapies.

## Results

### EZH2 inhibitors suppress the proliferation of AR-positive prostate cancer cells

To determine the gene(s) essential for the sustained proliferation of both androgen-dependent and castration resistant prostate cancer (CRPC) cells in an unbiased manner, we conducted CRISPR-Cas9 knockout screening in both parental, hormone-dependent LNCaP cells and its hormone-refractory counterpart LNCaP-abl (abl) cells under their respective proliferating conditions (Figure 1A). Positive control genes that are known to be required for cell proliferation in general were strongly selected, suggesting that the screens could reliably identify new essential genes (Figure S1A). We used MAGeCK (Chen et al., 2018; Li et al., 2015) to analyze the CRISPR screens, which assigns a beta score to each gene to approximate change of CRISPR guide RNA abundance. Therefore, in the cells grown for 4 weeks compared with those on day 0, more negative beta score represents higher dependency of cell growth on the target gene. We were particularly interested in genes that are more essential for the androgen-independent growth of CRPC cells, so we compared the beta scores between LNCaP and abl. Although most genes possessed similar beta scores between these two prostate cancer cell lines, we did find some showing significantly lower scores in abl than in LNCaP (Figure S1B). For example, several transcription factors that are well known for their prominent roles in CRPC, such as AR, FOXA1 and MYC, were consistently identified(Labbe and Brown, 2018). Interestingly, EZH2 was one of the top hits that exhibit much stronger dependency in abl cells than in LNCaP (Figure 1B), while EZH1, another mammalian homolog of Drosophila Enhancer of Zeste, was not required for either cell lines. This result is consistent with our previous report that genetic inhibition of EZH2 better suppressed growth of CRPC cells than that of androgen-dependent prostate cancer cells (Xu et al., 2012). Since EZH2 is a druggable enzyme with readily available small-molecule inhibitors, we assessed the efficacy of EZH2-targeting drugs in prostate cancer cells. We tested two compounds, GSK126 (McCabe et al., 2012b) and EPZ-6438 (Knutson et al., 2012), in a panel of human prostate cell lines, including two benign prostate epithelial cells, two androgen receptor (AR)-null prostate cancer cells, and eight AR-positive cancer cells. Only malignant cells with intact AR signaling, especially the hormone-refractory lines, were sensitive to both inhibitors (Figure 1C). Concentrations as low as 500 nM of the EZH2 inhibitors (EZH2i) greatly retarded the growth of castration-resistant abl cells (Figure 1D), and relatively higher doses of EZH2i were required to suppress the androgen-dependent LNCaP cells (Figure 1E). The inhibitory effect of EZH2i was minimal in AR-null DU145 (Figure S2A). This was consistent with a previous report that neither PC3 nor DU145 requires EZH2 for continued growth (Garapaty-Rao et al., 2013). Cell cycle analysis showed that EZH2i induced G0-G1 arrest in responsive CRPC cell lines within 3 days of the drug treatment (Figure S2B), but no cytostatic effect was observed in the unresponsive DU145 cells (Figure S2C). To further evaluate EZH2i effect on CRPC cell growth *in vivo*, we treated subcutaneous xenografts of the hormone refractory CWR22Rv1 cells in castrated mice with either GSK126 (Figure 1F) or EPZ-6438 (Figure 1G). Both compounds significantly retarded tumor growth following 21 days of treatment. Taken together, we demonstrated that EZH2 is a promising therapeutic target for prostate cancer and that EZH2 inhibitors may benefit patients with AR-positive, metastatic, hormone-refractory tumors.

**Figure 1.**
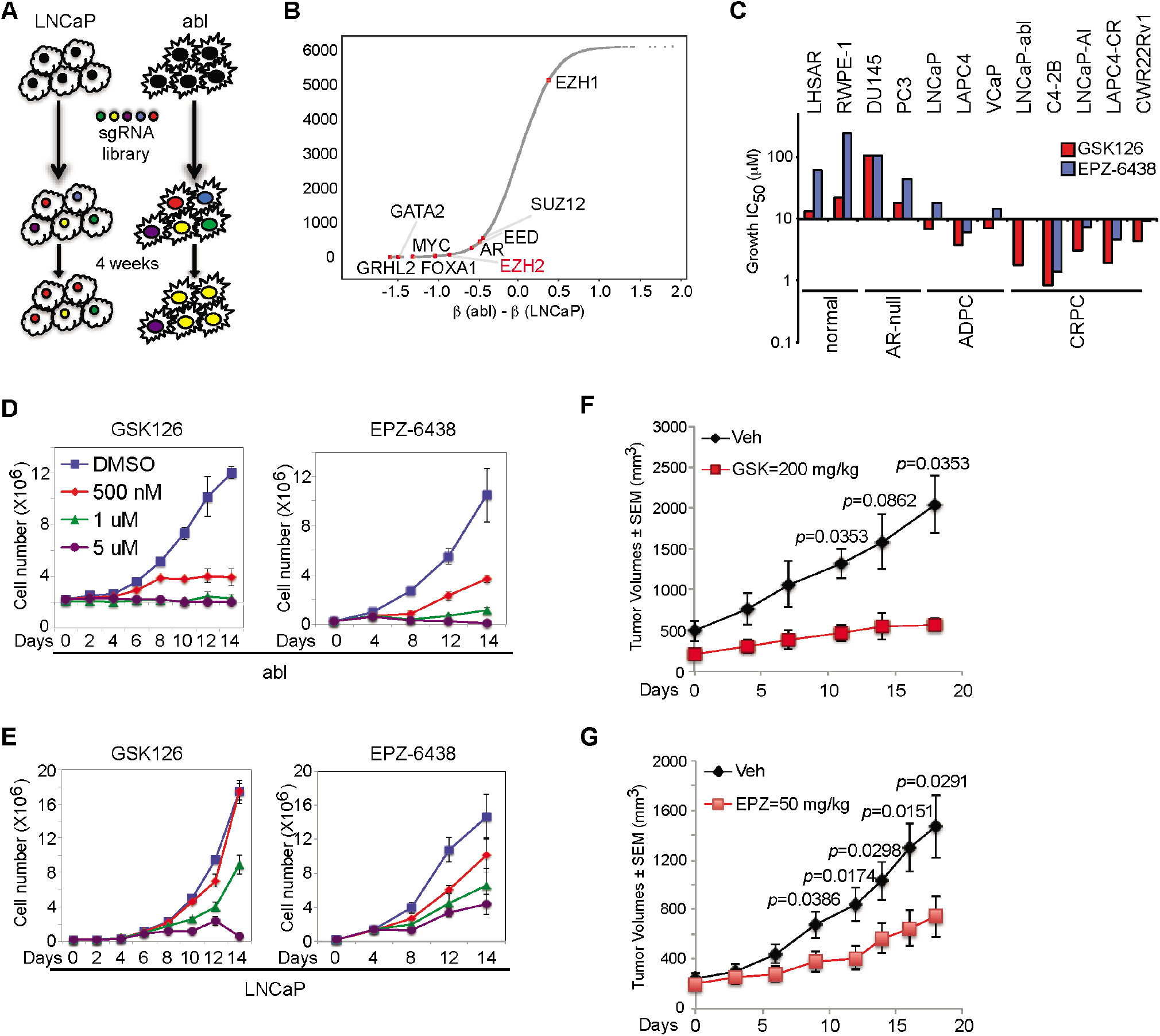
Inhibitors of EZH2 methyltransferase activity showed potent inhibitory effects in prostate cancer cells, especially the castration resistant ones. (A) Work flow of CRISPR knockout screens in LNCaP and abl cells. (B) Difference of gene essentialities, represented by beta (β) scores, between abl and LNCaP [β(abl)-β(LNCaP)] for each individual gene. Positions of representative genes were indicated by red dots. (C) IC50 for two EZH2 inhibitors (GSK126, red bars; EPZ-6438, blue bars) in a panel of prostate normal and cancer cell lines after 6 days of treatment. Cells were grouped based on the basic characteristics. ADPC, androgen-dependent prostate cancer; CRPC, castration-resistant prostate cancer. (D-E) Effects of EZH2 inhibitors (left panels, GSK126; right panels, EPZ-6438) on cell growth over time in abl (D) and LNCaP (E) with indicated concentrations of the compounds. (F-G) Growth curves of xenograft tumors in castrated nude mice injected with CWR22Rv1 cells receiving vehicle (Veh) or GSK126 (F), or EPZ-6438 (G), at the indicated doses.

### EZH2 inhibition induces specific gene signatures in sensitive CRPC cells

To investigate the mechanisms underlying the action of EZH2 inhibitors in sensitive prostate cancer cells, we profiled the gene expression pattern of abl cells upon the treatment with GSK126 or EPZ-6438 (Figure 2A). Both compounds induced very similar transcriptional changes, indicating likely on-target genetic effects. Interestingly, a large number of genes were significantly downregulated instead of being de-repressed, whereas in susceptible DLBCL cells EZH2 inhibitors induced a robust transcriptional activation(McCabe et al., 2012b). Genes that were commonly downregulated by both EZH2i in abl cells were significantly enriched in DNA damage response, such as various DNA damage repair (DDR) pathways, cell cycle regulation and DNA replication process (Figure 2B). There were no significant functional annotations for the upregulated genes in abl. To rule out the possible off-targeted effect of EZH2i on gene regulation, we carried out two analyses. First, we compared transcriptional profiles in abl being treated with EZH2i or EZH2-targeting RNA interference (RNAi) (Xu et al., 2012), and found they led to highly similar gene expression patterns (Figure 2C). Second, EZH2 inhibitors mainly de-repressed genes in EZH2i-insensitive DU145 cells (Figure S3A). These genes were consistently upregulated by EZH2 silencing (Figure S3B), recapitulating the suppressive function of EZH2 as an H3K27 methyltransferase. We validated several differentially expressed genes, and showed that EZH2i suppressed their expression in a dose- (Figure S4A) and time-dependent manner (Figure S4B). We further examined the expression of these genes in three other EZH2i-sensitive CRPC cell lines: C4-2B, LAPC4-CR and LNCaP-AI (Figure 2D). While GSK126 and EPZ-6438 upregulated EZH2-repressed genes to some extent, EZH2i decreased expression of EZH2-activated genes more uniformly in these cells. We further validated these findings in CWR22Rv1 xenografts, demonstrating consistent decrease in mRNA (Figure 2E) and protein (Figure 2F) levels of EZH2-activated genes by either compound. Taken together, these results indicated that EZH2 inhibitors indeed target the methyltransferase activity of EZH2, and yet they downregulate a group of EZH2-activated genes in CRPC cells.

**Figure 2.**
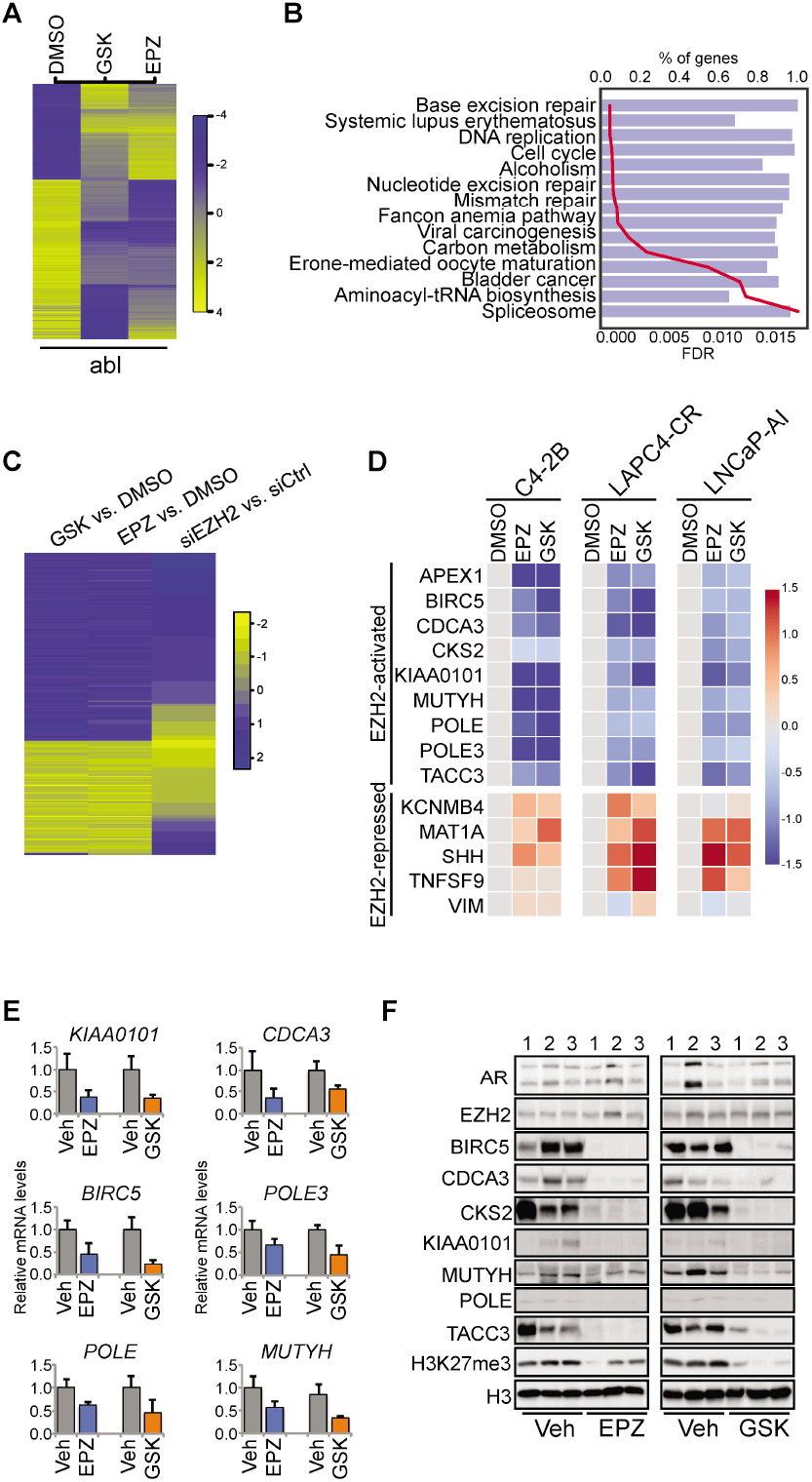
EZH2 inhibitors downregulated a large number of genes in sensitive castration resistant prostate cancer cells. (A) Heat map of differential gene expression patterns in abl cells being treated with vehicle (DMSO), 5 uM GSK126 (GSK) or 5 uM EPZ-6438 (EPZ) for 72 hrs. (B) Top overrepresented functional annotations of genes that were significantly downregulated upon the treatment with EZH2 inhibitors in abl cells from Gene Set Enrichment Analysis. Blue bars, percentage of genes in each specific functional category; red line, values of the false discovery rate (FDR) for the particular GO term. (C) Heat map of differential genes induced by EZH2 inhibitors (GSK, GSK126; EPZ, EPZ-6438) comparing to vehicle control (GSK vs. DMSO and EPZ vs. DMSO) or EZH2 knockdown comparing to control siRNA (siEZH2 vs. siCtrl) in abl cells. Gene expression profiling upon silencing of EZH2 was retrieved from our prior work(Xu et al., 2012). (D) Heat maps of quantitative real-time RT-qPCR results showing changes in mRNA levels of selected gene in three prostate cancer cell lines (C4-2B, LAPC4-CR and LNCaP-AI). Cells were treated with vehicle (DMSO), 5 uM GSK126 (GSK) or 5 uM EPZ-6438 (EPZ) for 72 hrs. Top panels, EZH2-activated genes; bottom panels, EZH2-repressed genes. (E-F) Evaluation of mRNA expression (E) or protein levels (F) of selected genes in xenograft tumor tissues from control mice (Veh) or mice treated with GSK126 (GSK) or EPZ-6438 (EPZ). Numbers, triplicates of samples from control or treatment group.

### EZH2i-resistance mutations rescue the effects of EZH2 inhibitors on gene expression and cell growth

Several secondary mutations in EZH2, including Y111D and Y661D, were identified in DLBCL clones that became refractory to EZH2i (Baker et al., 2015; Bisserier and Wajapeyee, 2018). These resistance mutants regain methyltransferase activity in the presence of EZH2i, and thus offer a genetic means to evaluate the targeted action of EZH2i. We replaced endogenous EZH2 in abl with these mutants, and then evaluated the responses of both parental and mutant cell lines to EZH2i by assessing H3K27 methylation, cell proliferation and gene expression. When either Y111D or Y661D was expressed, EZH2i-induced reduction of H3K27me3 level was dramatically alleviated (Figure 3A). This was especially notable in the case of Y111D, which completely abolished the effect of EZH2i on H3K27me3. In cell growth assays, EZH2i failed to suppress the growth of abl cells expressing Y111D or Y661D (Figure 3B, Figure S5A and S5B), even after incubation with the drugs for up to 15 days (Figure 3C). These results demonstrated that EZH2 mutations that render EZH2i resistance in DLBCL also confer similar refractory phenotype in prostate cancer cells. Intriguingly, both Y111D and Y661D rescued the expression of EZH2-activated genes in the presence of EZH2i (Figure 3D). Interestingly, this rescue was less evident for EZH2-repressed genes (Figure S5C). These findings provide strong support for the conclusion that EZH2 inhibitors abrogated prostate cancer cell growth through specific blockade of EZH2 functions. They also support that genes transactivated by EZH2 might predominantly mediate the action of EZH2 inhibitors in CRPC.

**Figure 3.**
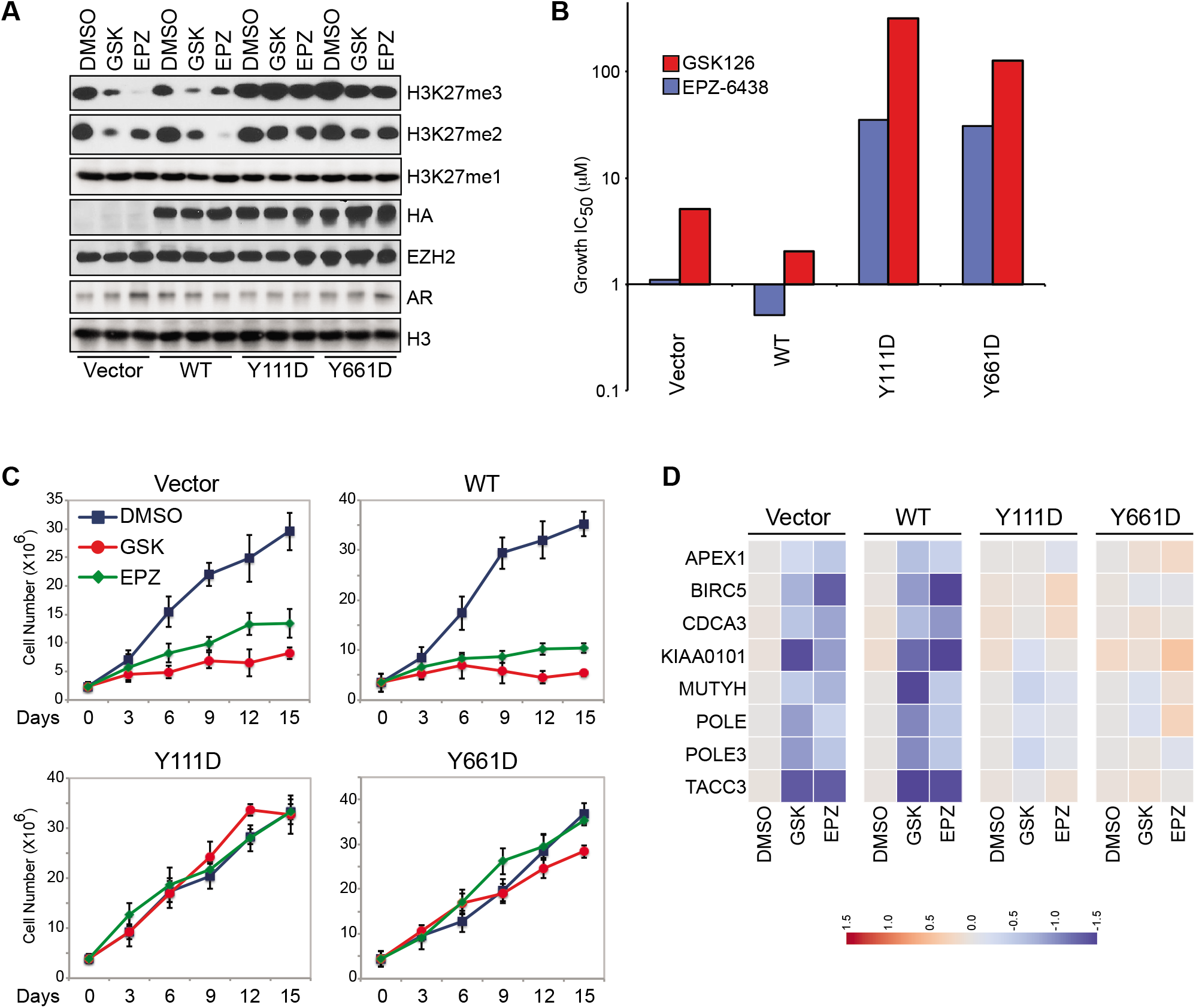
Genetic mutations of EZH2 conferred resistance to the effects of EZH2 inhibitors on cell growth and gene expression. (A) Western blots of H3K27 methylation levels in abl cells that were replaced with the control (Vector), the wild type (WT) or EZH2 mutants bearing different point mutations (Y111D or Y661D). Cells were treated with DMSO or EZH2 inhibitors (GSK, GSK126; EPZ, EPZ-6438) for 72 hrs. (B) IC50 values for two EZH2 inhibitors in abl cells expressing the control (Vector), the wild-type EZH2 (WT) or the indicated mutants. Cells were incubated with EZH2 inhibitors (GSK126, red bars; EPZ-6438, blue bars) for 6 days and then collected for direct counting after trypan blue staining. (C) Effects of EZH2 inhibitors (GSK, GSK126; EPZ, EPZ-6438) on the growth of abl cells expressing the control (Vector), the wild-type EZH2 (WT) or the mutants at indicated time points. (D) Expression of EZH2-activated genes was detected by RT-qPCR in abl cells that were replaced with the control (Vector), the wild-type EZH2 (WT) or the mutants in the presence of vehicle (DMSO) or 5 uM EZH2 inhibitors (GSK, GSK126; EPZ, EPZ-6438) for 3 days.

### EZH2 inhibitors decrease global H3K27me3 signal on chromatin regardless of cellular response to the compounds

In view of the canonical function of EZH2 in catalyzing H3K27me3, we set out to investigate how the repressive chromatin mark contributed to the growth-inhibitory and gene-regulatory effects of EZH2i. All of the tested prostate cell lines demonstrated reduced total H3K27 di- and tri-methylation levels in a dose-dependent manner with either compound (Figure 4A), which is in contrast with their distinct cellular growth responses to EZH2i. Discrepancies between EZH2i-induced H3K27me3 reduction and cell insensitivity to EZH2i were also reported in lymphoma cells (McCabe et al., 2012b; Qi et al., 2012). To accurately evaluate the locus-specific changes of H3K27me3 on chromatin, we adopted the ChIP-Rx method, which uses a “SPIKE-IN” strategy to quantify genome-wide histone modification relative to a reference epigenome with defined quantities (Orlando et al., 2014). Canonical normalization methods such as using the total sequencing reads showed moderate H3K27me3 changes after cells were treated with EZH2i (Figure S6A), while normalization to the reference epigenome showed pronounced H3K27me3 reductions (Figure 4B). In the EZH2i-insensitive DU145 cells, we noticed a similar or even more robust EZH2i-induced decrease of H3K27me3 intensity than in abl (Figure 4C and Figure S6B). This implies that reduction of H3K27me3 alone does not confer growth sensitivity to EZH2i. Indeed, we detected a moderate decrease in overall H3K27me3 amount and its signal on chromatin in abl cells as early as 2 days of EZH2i treatment (Figure 4D), and DU145 cells showed a marked loss of the histone modification within the same time frame (Figure S6C). To find any functional significance of EZH2i-triggered H3K27me3 alterations in abl cells, we associated changes of the repressive histone mark with differential gene expression upon compound treatment (Figure 4E). Although basal level of H3K27me3 was noticeably higher at the promoter regions of EZH2i-upregulated genes, there were no differences regarding the extent of H3K27me3 decrease among EZH2-repressed, EZH2-activated or EZH2-indifferent genes. This result suggests that while the steady status of H3K27me3 is associate the silenced transcription of downstream targets, the fluctuation of H3K27me3 signals does not always lead to immediate transcriptional changes of nearby genes. Taken together, these results suggest that H3K27 methylation, a readout of the polycomb repressive function of EZH2, may not be the determining factor of transcriptional changes or cellular responses to EZH2i.

**Figure 4.**
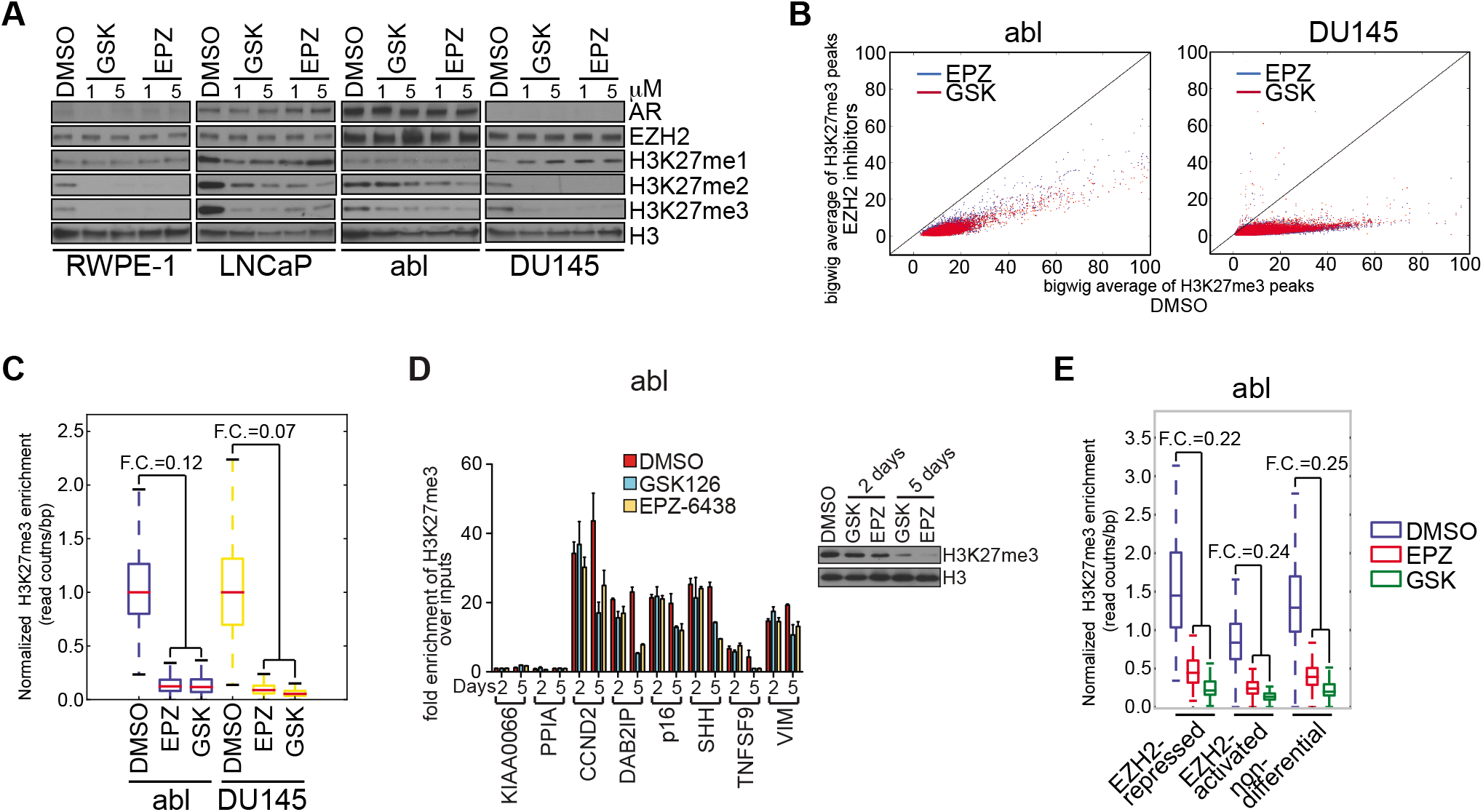
Genome-wide reduction in H3K27 trimethylation levels alone did not dictate the action of EZH2 inhibitors in prostate cancer cells. (A) Evaluation of H3K27 methylation levels in prostate cell lines with the treatment of vehicle (DMSO), GSK126 (GSK) or EPZ-6438 (EPZ) at specified doses (1 or 5 μM final concentration). (B) Scatter plots of H3K27me3 peak signals, after being normalized to the Drosophila reference epigenome, in abl and DU145 cells under control condition (x-axis) or after the treatment with EZH2 inhibitors (y-axis). (C) Comparison of EZH2 inhibitor-induced changes in H3K27me3 levels between abl and DU145 cells after SPIKE-IN normalization. GSK, GSK126; EPZ, EPZ-6438; F.C., fold change. (D) Direct ChIP-qPCR of H3K27me3 at selected chromatin regions after abl cells were treated with control (DMSO) or EZH2 inhibitors (GSK126 or EPZ-6438) for indicated number of days. KIAA0066 and PPIA, negative controls; right panel, H3K27me3 protein levels by immunoblotting in the corresponding ChIP samples. GSK, GSK126; EPZ, EPZ-6438. (E) SPIKE-IN normalized signals of H3K27me3 peaks around genes that were upregulated (EZH2-repressed), downregulated (EZH2-activated) or showed no differences (non-differential) upon the treatment with EZH2 inhibitors. Intensities of the histone mark under either control (DMSO) or treatment condition (GSK, GSK126; EPZ, EPZ-6438) were plotted. F.C., fold change.

### A core gene signature in regulation of DNA damage repair is essential for the growth-inhibitory effects of EZH2 inhibitors in prostate cancer

We next sought to investigate the molecules underlying the inhibitory effects of EZH2i. To this end, we conducted CRISPR-Cas9 knockout screens in abl with or without GSK126 treatment (Figure 5A). As a control, we performed the CRISPR screens in LNCaP in parallel. Most genes showed similar beta scores using MAGeCK between treatment and control conditions in the two cell lines (Figure S7A). To search for genes that mediate specifically the biological effects of GSK126 in CRPC, we defined Δβ as the difference in beta scores between treatment (+GSK126) and control (+vehicle) groups. We postulated that knockout of genes crucial for the growth-inhibitory effect of EZH2i would render clones resistant to the compounds. Therefore, genes with positive Δβ (i.e., less essentiality in treatment condition than control condition) are required for EZH2i activity. This analysis identified a group of genes specifically in abl cells, majority of which function in DNA damage repair (Figure 5B). However, these genes were not selected from the CRISPR-Cas9 screening in LNCaP (Figure S7B). Gene set enrichment analysis (GSEA) of genes with positive Δβ values in abl confirmed the significant enrichment of DNA repair processes, especially the base excision repair (BER) pathway (Figure 5C), while similar analyses of genes with positive Δβ values in LNCaP did not find functional enrichment (Figure S7C). Thus, CRISPR screens suggest that DNA repair genes are directly targeted by EZH2i and therefore indispensible for the biological effects of these compounds in prostate cancer cells.

**Figure 5.**
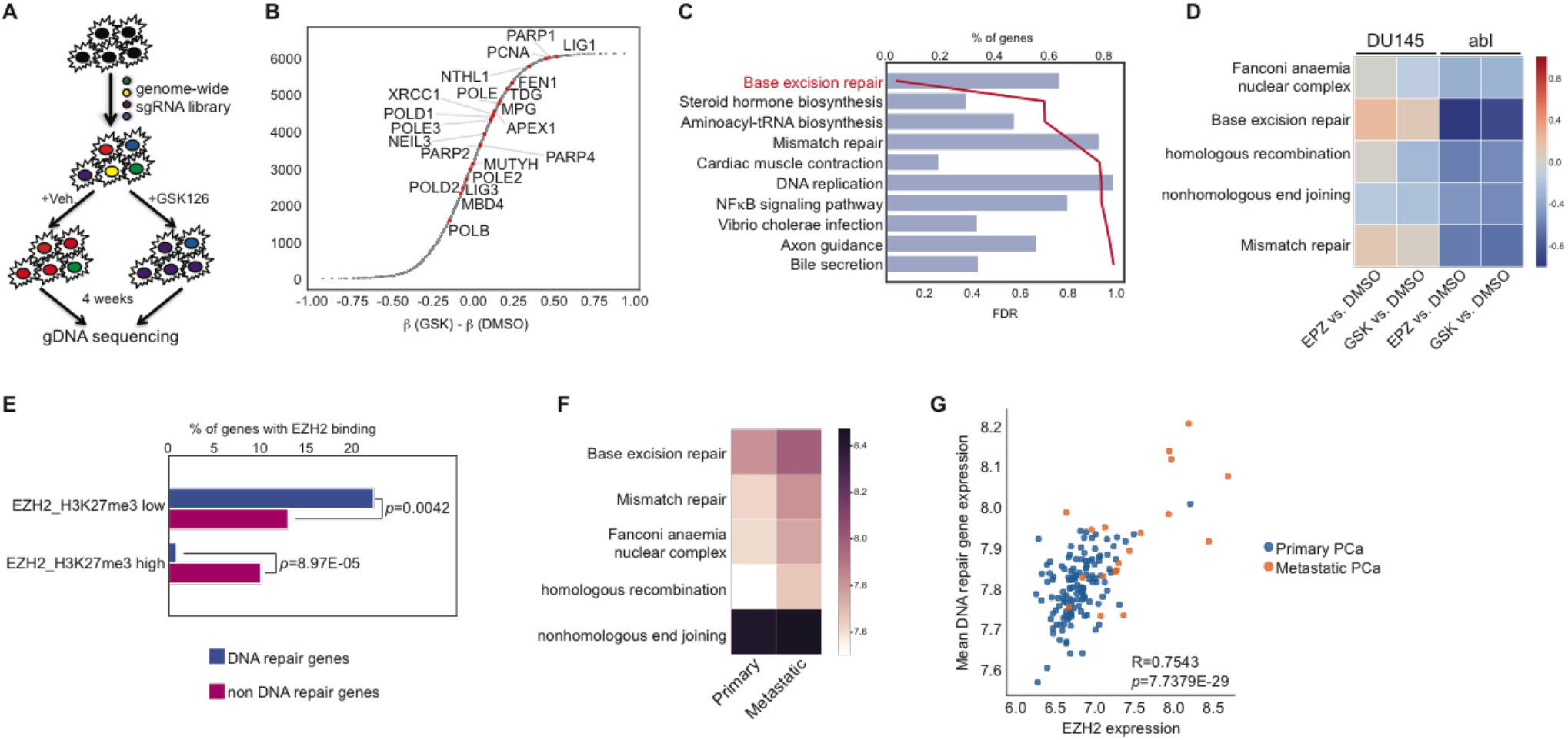
DNA damage repair was crucial for the biological effects of EZH2 inhibitors in prostate cancer. (A) Work flow of CRISPR-Cas9 knockout screening, which targets 6,000 cancer-related genes in LNCaP and abl cells in the presence of vehicle or GSK126 (EZH2 inhibitor) for four weeks. (B) Distribution of delta beta (Δβ) scores, defined as beta scores under treatment (+GSK126) condition minus beta scores under control (+Veh.) condition [β(GSK)-β(DMSO)]. Representative DNA repair genes were indicated by red dots. (C) Gene set enrichment analysis of genes with positive delta beta scores (Δβ) in CRISPR-Cas9 knockout screening in abl cells. Blue bars, percentage of genes in each specific functional category; red line, values of the false discovery rate (FDR) for the particular gene ontology term. (D) Heat map showing differential expression (log-transformed) of genes involved in the specified DNA damage repair pathways upon treatment with EZH2 inhibitors. Transcript levels of these genes in abl and DU145 cells were compared between treatment groups and control group (EPZ vs. DMSO and GSK vs. DMSO). (E) Percentages of genes containing two types of EZH2 chromatin binding (EZH2_H3K27me3 low, EZH2 binding sites with low or no H3K27me3 signals; EZH2_H3K27me3 high, EZH2 binding sites with high H3K27me3 signals) within 1kb around their transcriptional start sites. Blue bars, genes involved in DNA damage repair pathways; rose red bars, genes not involved in DNA damage repair. (F) Heat map showing expression of DNA repair genes in primary or metastatic prostate cancer (Taylor et al., 2010). Genes were categorized into different DNA damage repair pathways according to their functions. (G) Expression correlation between EZH2 and DNA repair genes in a prostate cancer cohort (Taylor et al., 2010). Each dot represents a clinical sample, with primary prostate cancer (PCa) samples colored in blue while the metastatic ones colored in yellow.

Indeed, EZH2i treatment significantly downregulated expressions of multiple DNA damage repair pathways in abl, but not in DU145 (Figure 5D). These genes represent direct targets of EZH2, as EZH2 binding is significantly enriched within ±1 kb of transcription start sites (TSSs) of these genes (Figure 5E). Interestingly, this binding enrichment was only observed for EZH2 binding peaks with low or absent H3K27me3 enrichment, whereas H3K27me3-associated EZH2 peaks were excluded from the promoter regions near these genes. This is in line with the conclusion that H3K27me3 is irrelevant to the activity of EZH2i in CRPC. To further validate the clinical significance of DNA damage repair pathways, we retrieved their transcript levels in two independent prostate cancer cohorts (Grasso et al., 2012; Taylor et al., 2010), and found that they were significantly elevated in metastatic CRPC compared to primary prostate tumors (Figure 5F and Figure S7D). In addition, expression of these genes shows very strong positive correlation with that of EZH2, both displaying much higher levels in metastatic advanced prostate cancer (Figure 5G and Figure S7E). Moreover, expression of these genes in prostate cancer cells, especially genes in BER and mismatch repair pathways, is correlated with the cellular sensitivity to EZH2i, which is the lowest in DU145, intermediate in LNCaP, and highest in abl (Figure S7F). Taken together, our findings revealed a close connection between the tumor-suppressive activity of EZH2 inhibitors and DNA repair machinery, which suggests a new indication for these compounds as anticancer drugs.

### EZH2 inhibitors enhance responses of prostate cancer cells to DNA damaging agents

Our data above provides a rationale for applying EZH2i to sensitize CRPC cells to DNA damage. To validate this, we exposed abl, LNCaP and DU145 cells to increasing doses of ionizing radiation (IR). While abl cells became much more sensitive to IR after GSK126 treatment (Figure 6A), this difference was very minor in LNCaP (Figure 6B) and non-existent in DU145 (Figure 6C). In addition, when 5 gray (Gy) of IR was applied and cell recovery was monitored, we observed drastically delayed DNA damage repair (DDR) in abl pretreated with GSK126 (Figure S8A), but neither in LNCaP (Figure S8B) nor in DU145 (Figure S8C). It is worth noting that without EZH2i pretreatment, IR has moderate effects in abl even at a dose as high as 20 Gy (Figure S8D). This suggests that abl may represent a radioresistant scenario and EZH2 inhibitors may be considered to overcome radiotherapy resistance in advanced prostate cancer. PARP-1 is an ADP-ribosylating enzyme involved in various forms of DNA repair, including BER (Ko and Ren, 2012). It has been reported that deficiencies of any components in BER pathway resulted in hypersensitivity of cancer cells to PARP inhibitors (Horton et al., 2014). We therefore assessed the biological effect of combining EZH2i with the PARP-1 inhibitor, olaparib, on proliferation of LNCaP or abl cells. Combined treatment of EZH2i and olaparib greatly suppressed abl cell growth compared to each drug alone (Figure 6D), and showed strong synergistic effect (Figure 6E, right panel). However, this synergy of EZH2i and olaparib was much milder in LNCaP (Figure 6E, left panel). Taken together, our work has uncovered a novel therapeutic strategy for hormone-independent prostate cancer, which exploits the suppressive effects of EZH2 inhibitors on DNA damage repair genes to sensitize cancer cells to DNA damaging agents.

**Figure 6.**
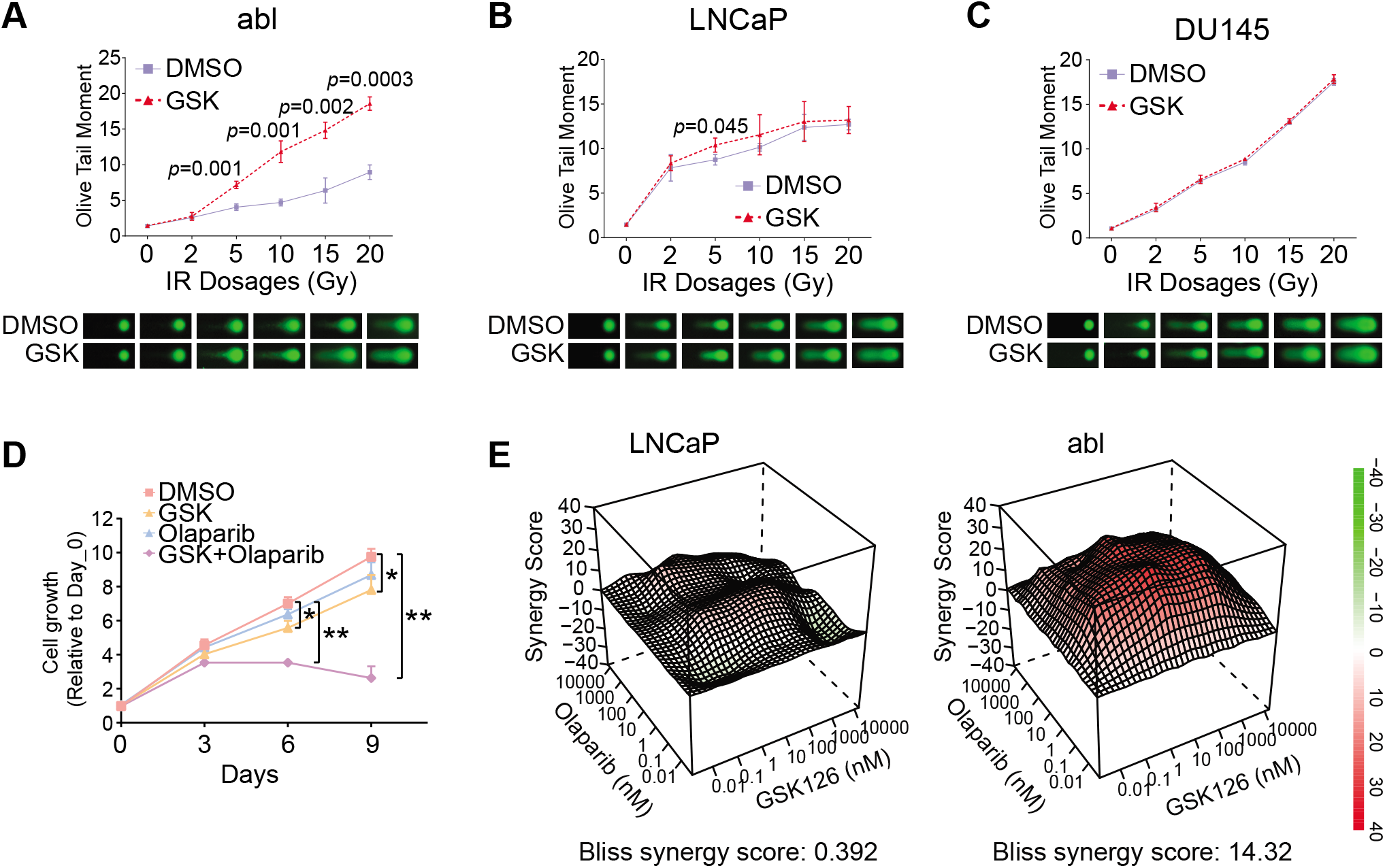
EZH2 inhibitors enhanced sensitivity of prostate cancer cells to DNA damage agents. (A-C) Results of the alkaline comet assays in abl (A), LNCaP (B) or DU145 (C). Prostate cancer cells were pretreated with vehicle (DMSO) or 5 μM GSK126 (GSK) for 7 days, and then exposed to increasing dosages of ionizing radiation (IR) followed by recovery for 4 hrs. Top panels, quantification of the results using “Olive Tail Moment” parameter; bottom panels, representative images of each IR dosage. (D) Combination therapy of EZH2 inhibitor and olaparib to abrogate the androgen-independent growth of abl cells. Cells were treated with vehicle (DMSO), 0.5 μM GSK126 (GSK) alone, 1 μM Olaparib alone, or both drugs (GSK + Olaparib) for indicated days, and cell proliferation were determined using Cell Counting Kit-8. *, *p*<0.05; **, *p*<0.01. (E) Synergistic effect of EZH2 inhibitor and olaparib in LNCaP (left panel) and abl (right panel) cells. Various doses of GSK126 and olaparib were combined and applied for 7 days. A matrix for synergy score was calculated and indicated in y-axis (Ianevski et al., 2017).

### Expression of DNA damage repair genes predicts sensitivity of cancer cells expressing wild-type EZH2 to EZH2 inhibitors

We next sought to determine whether what we observed in prostate cancer cells could be generalized to other types of wild-type EZH2-expressing solid tumors. We excluded diffuse large B-cell lymphoma (DLBCL) cells, acute myeloid leukemia (AML) cells, and multiple myeloma (MM) cells, as these types of hematopoietic malignancies often contain gain- or loss-of function mutations of EZH2 (Cancer Genome Atlas Research et al., 2013a; Makishima et al., 2010; Morin et al., 2010; Morin et al., 2011). Mutant EZH2 in these scenarios has been reported to exert its oncogenic function through a classical PRC2-dependent mechanism (Morin et al., 2010; Saygin et al., 2018). We also excluded solid tumor cell lines with genetic alterations of EZH2.

We first explored the expression correlation between wild-type EZH2 and the DNA damage repair genes in solid tumor cell lines using the Cancer Cell Line Encyclopedia (CCLE) data set (Barretina et al., 2012). Expression of DNA damage repair genes and EZH2 was strongly correlated (Figure 7A). We further confirmed this finding using clinical samples from the TCGA data sets (Cancer Genome Atlas Research et al., 2013b), and found that a strong positive expression correlation still holds true even within individual solid tumor types (Figure 7B and S9A). These results suggest that EZH2-mediated control of DNA damage repair machinery may represent a common mechanism of EZH2 oncogenic function in solid tumors without EZH2 mutations. To examine whether expression of DNA damage repair genes can dictate EZH2i sensitivities, we analyzed Cancer Therapeutics Response Portal (CTRP) compound screening data (Rees et al., 2016), which measured sensitivities to the EZH2 inhibitor BRD-K62801835-001-01-0 (BRD) in 531 wide-type EZH2-expressing cancer cells lines. When comparing across distinct tissue types, cellular sensitivity to EZH2 inhibitor is significantly correlated with expression of DNA repair genes in the positive direction (Figure 7C). In comparison with the bottom one third of cell lines with the lowest levels of DNA damage repair genes, the top one third that express the highest levels of these genes were much more susceptible to EZH2 inhibitor treatment (Figure S9B). In contrast, genes that were repressed by EZH2, although having negative expression correlation with EZH2 (Figure S9C), do not consistently predict EZH2i sensitivity in solid tumor cells without EZH2 mutations. No matter if it is across all cell lines (Figure S9D) or among individual tissue types (Figure S9E), higher levels of EZH2-repressed genes are not associated with less sensitivity to EZH2i. Furthermore, we confirmed that expression of DNA repair genes predicts EZH2 dependency using genome-wide CRISPR-Cas9 knockout screen data across 489 cell lines harboring no EZH2 mutations (Tsherniak et al., 2017). Cancer types with higher expression of DNA repair genes are more dependent on EZH2 for sustained growth, reflected by more negative CERES scores for EZH2 (Figure 7D). Taken together, our analyses suggest that the mechanism of inhibitory action of EZH2 inhibitors may be different in solid tumors expressing wild-type EZH2 from that in hematopoietic malignancies with EZH2 somatic mutations. The group of DNA damage repair genes that are activated by EZH2 can reliably predict efficacy of EZH2 inhibitors in wild-type EZH2-expressing solid tumors.

**Figure 7.**
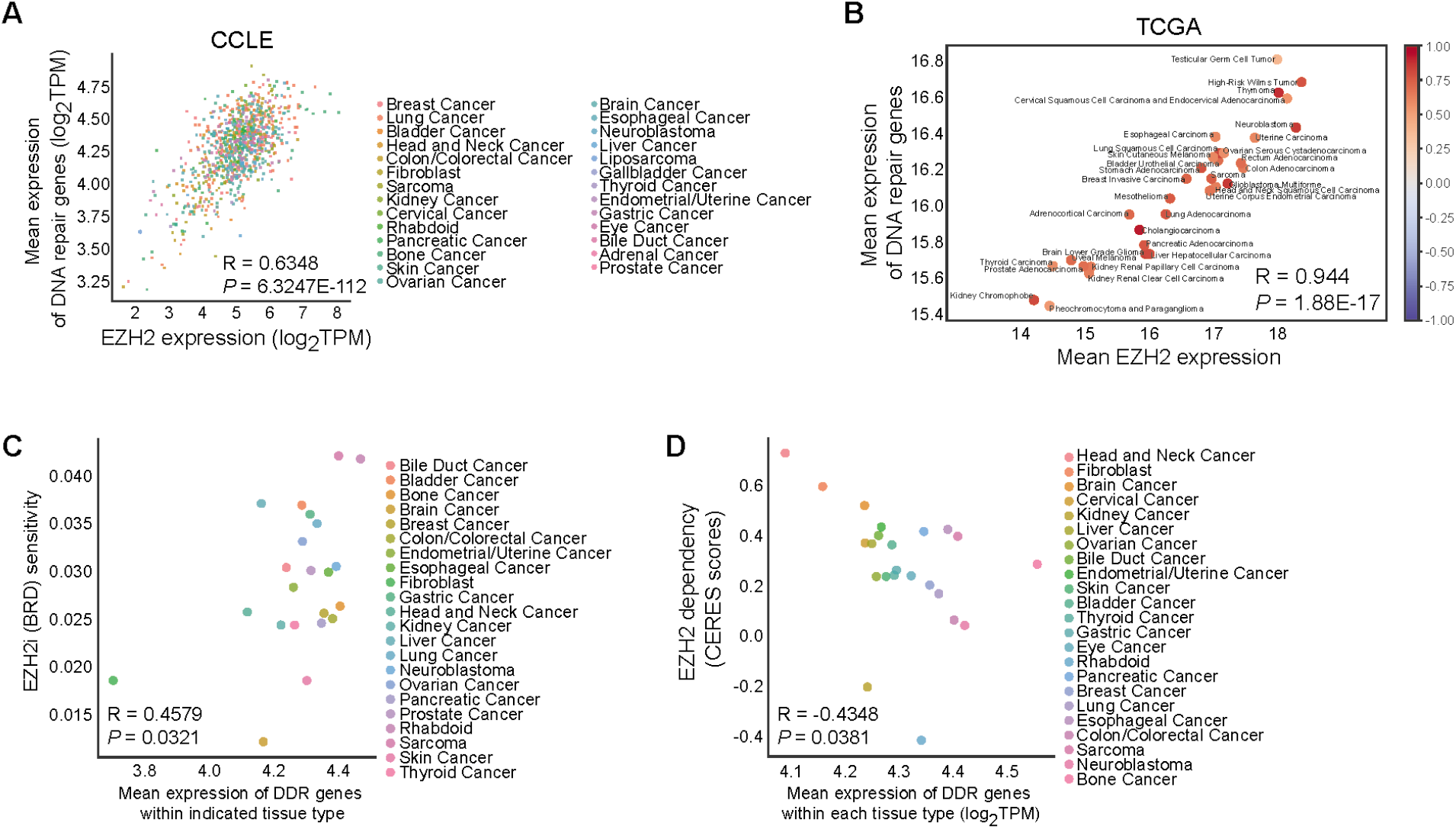
Levels of DNA damage repair genes predict cellular responses to EZH2 inhibitors in multiple types of solid tumors that contain no mutations of EZH2. (A-B) Expression correlation between EZH2 and DNA repair genes in cancer cells from CCLE gene expression data (Barretina et al., 2012) (A) or in patient samples from TCGA clinical data (Cancer Genome Atlas Research et al., 2013b) (B). Each dot represents one cell line that was illustrated with the same color to indicate the same cell type (A), or represents one cancer type with their names by side (B). (C-D) Dot plots demonstrating the association between expression of DNA damage repair (DDR) genes and sensitivity to EZH2 inhibitor (C) or EZH2 dependency (D) among indicated cell types. Cellular sensitivity to EZH2 inhibitor (BRD, BRD-K62801835-001-01-0) and EZH2 dependency scores (CERES scores) in wild-type EZH2-expressing solid tumor cell lines were derived from CTRP compound screen data (Rees et al., 2016) and DepMap CRISPR-Cas9 knockout screening data (Tsherniak et al., 2017), respectively. Individual cell types were highlighted in different colors and specified.

## Discussion

The methyltransferase EZH2 has been a focus of cancer drug development for several years. Inhibitors of EZH2 have been tested in patients with non-Hodgkin lymphoma harboring gain-of-function (GOF) mutations of EZH2 (Gulati et al., 2018). This therapeutic strategy is based on the concept that these mutations lead to increased levels of H3K27me3 and robust silencing of tumor suppressor genes. Therefore, treatment with EZH2 inhibitors leads to reactivation of these tumor suppressor genes and growth inhibition in these cases. A wide range of solid tumors including prostate cancer expresses high levels of wild-type EZH2, and therefore whether EZH2 inhibitors will have activity in solid tumors expressing only wild-type EZH2 remains an open question. Our study in prostate cancer cells revealed that distinct from its role in PRC2-mediated transcriptional repression, EZH2 directly activates a set of DNA damage repair genes. A gene signature based on these EZH2-activated genes underlies the growth-suppressive effects of EZH2 inhibitors in CRPC cells and predicts response to EZH2i across cancer types lacking EZH2 mutations. Our findings define a potentially new mechanism of action of EZH2 inhibitors, and lay the mechanistic foundation for the potential clinical applications of these compounds to sensitize cancer cells to DNA damaging agents.

CRISPR-based knockout screens revealed that DNA damage repair genes, particularly those in the BER pathway, are required for the optimal growth inhibitory effects of EZH2i in CRPC cells. The expression of these genes was acutely and robustly downregulated upon pharmacological inhibition of EZH2, and their levels are highly correlated with cellular sensitivities to EZH2 inhibitors. These findings suggest a non-canonical mechanism of action of these drugs in solid tumors that harbor no genetic alterations of EZH2 and have an important impact on clinical development of these drugs in these scenarios. The lack of EZH2 mutations in common solid tumors including prostate cancer raised concerns that conducting trials of EZH2 inhibitors in these cancers would be ineffective. In this study, we demonstrated that EZH2 inhibitors could be efficacious in wild-type EZH2 expressing solid tumors, such as CRPC. This effectiveness is positively associated with the expression of DNA damage repair genes, which can potentially be used to predict EZH2i sensitivities in solid tumors that contain no EZH2 mutations. This is in contrast to the NHL cases. In lymphomas, such as DLBCL and follicular lymphoma (FL), cases with activating somatic mutations of EZH2 are generally more susceptible to EZH2 inhibitors (Tremblay-LeMay et al., 2018), and presence of these genetic alterations is predictive of EZH2i efficacy. However, there exist some wild-type EZH2 expressing DLBCL cells that are also sensitive to EZH2 inhibitors, and the DNA repair genes we identified in this study may possibly account for the inhibitory effects of the drugs in these cases. Therefore, we suggest that the EZH2-induced gene signature defined in our study may be useful as a predictive biomarker for response to EZH2 inhibitors in cancers lacking GOF mutations.

Another important implication of our findings is the potential of a new combination therapy strategy leveraging the interaction between EZH2 and the DNA damage repair machinery. Pretreatment with EZH2 inhibitors significantly boost the growth inhibitory effects of DNA damaging agents, such as ionizing radiation and olaparib. Thus, combination of EZH2 inhibitors with genotoxic agents may be a tenable approach in anticancer therapy. Evidence from other studies supports our model. Treatment with GSK126 in chemotherapy resistant B-cell lymphoma cells led to downregulation of MDMX, the negative regulator of p53. Subsequently, cells became re-sensitized to etoposide-induced apoptosis (Smonskey et al., 2016). EZH2 inhibition has also been demonstrated to dramatically enhance the efficacy of olaparib in BRCA-deficient breast cancer cells (Caruso et al., 2018; Yamaguchi et al., 2018). In this case, EZH2 was found to be PARylated by PARP1, which led to reduced H3K27me3. Furthermore, the modified EZH2 was detected at sites of DNA damage and controlled DNA repair in an H3K27me3-independent manner. Our result that EZH2 specifically activates a set of DNA repair genes provides an explanation for the previously unknown mechanism underlying EZH2-regulated DNA damage repair. Interestingly, the LNCaP-abl CRPC cell line in which we demonstrated the synergistic effects of EZH2 inhibitors and DNA damaging agents carries a heterozygous deletion of *BRCA2* exon 12 (Rauh-Adelmann et al., 2000) and is intrinsically refractory to genotoxic insults. Based on our data in CRPC cell models and our analysis across hundreds of cancer cell lines, it is plausible that EZH2 inhibitors may overcome resistance to DNA damaging agents in advanced cancer and improve the efficacy of olaparib in BRCA-deficient and -proficient tumors. Overall, these findings support the development of combination therapies that include DNA damaging agents and EZH2 inhibitors across a range of cancer types.

Our study has clearly illustrated a close link between DNA damage repair genes and the oncogenic function of EZH2. However, it remains unclear how EZH2, which lacks DNA binding domain and possesses H3K27 methyltransferase activity, activates the expression of DNA repair genes. In prostate cancer, several studies have demonstrated an involvement of AR in maintaining DNA repair efficiency (Goodwin et al., 2013; Polkinghorn et al., 2013). AR binds to the *cis*-regulatory elements of multiple DNA repair genes, and the resulting enhanced DNA repair capacity confers radioresistance. We have previously reported that EZH2 can serve as a co-activator of AR (Xu et al., 2012), it is plausible that EZH2 targets DNA damage repair genes through AR in prostate cancer. This mechanism, however, cannot explain the strong associations of EZH2 and DNA repair genes in various cancer types that do not express AR. Widely expressed transcription factors including PARP1/2 (Campbell et al., 2013) and E2F1 (Xu et al., 2016), have also been found to cooperate with EZH2 in gene regulation. These factors are known to be required for proper DNA damage responses (Biswas and Johnson, 2012; Javle and Curtin, 2011). Therefore, characterization of EZH2-centered proteomic network, especially at sites of DNA breaks, may shed light on the mechanism underlying EZH2-mediated regulation of the DNA repair pathway.

In summary, our study elucidated a novel mechanism of EZH2 inhibitor action in cancer. We identified a core gene signature involving DNA damage repair as a pharmacological readout of EZH2 inhibitor function. These DNA damage repair genes are directly targeted by EZH2 inhibitors and underlie the anti-tumor effects of these compounds. Finally, our data suggests that EZH2 inhibition might be an attractive approach to sensitize cancers that overexpress EZH2-regulated DNA damage repair genes to genotoxic agents.

## Experimental Procedures

### Antibodies and Reagents

Antibodies used in this study include: αAR (H-280, sc-13062) for ChIP-qPCR and ChIP-Seq; αAR (N-20, sc-816) for Western blot and immunoprecipitation; αH3K27me3 (C36B11, #9733S) for ChIP-qPCR, ChIP-Rx and Western blot; αH2Av (39715) for ChIP-Rx; αAR (441, sc-7305), αH3K27me1 (ab194688), αH3K27me2 (ab24684), αH3 (ab4086), αEZH2 (clone 11, 612666), αCDCA3 (FL-268, sc-134625), αKIAA0101 (SAB1406878), αTACC3 (C-2, sc-376883), αHA (51064-2-AP), αBIRC5 (D-8, sc-17779), αCKS2 (F-12, sc-376663), αMUTYH (C-6, sc-374571), αDNA pol ε A (34, sc-135885) for immunoblotting. EZH2 inhibitors were purchased from Xcessbio Biosciences Inc. (GSK126, M60071 and EPZ-6438, M60122) and olaparib (AZD2281) from Selleck Chemicals (S1060). The SMARTpool siRNAs (Dharmacon) used in this study were: siGENOME Non-Targeting siRNA Pool #2 (D-001206-14), SMARTpool ON-TARGETplus EZH2 siRNA (L-004218-00) and SMARTpool siGENOME EZH2 siRNA (M-004218-03).

### Normal and Cancer Prostate Epithelial Cell Lines and Culture Conditions

Benign and malignant prostatic epithelial cell lines RWPE-1, DU145, PC3, and LNCaP were originally purchased from the American Type Culture Collection. LHSAR cell line was kindly provided by Dr. Matthew Freedman. LAPC4, LNCaP-AI and LAPC4-CR were all obtained from Dr. Philip W. Kantoff’s lab. LNCaP-abl (abl) cell line was generously shared by Zoran Culig (Innsbruck Medical University, Austria). VCaP and CWR22Rv1 cell lines were graciously provided by Dr. Steven P. Balk. C4-2B was obtained from ViroMed Laboratories (Minneapolis, MN). All of these cell lines were authenticated at Bio-Synthesis Inc. and confirmed to be mycoplasma-free using MycoAlert Mycoplasma Detection Kit (Lonza). The specific culture conditions for each cell line were listed in Supplementary Table 1.

### Cell Proliferation Assay

Normal prostate epithelial cells and prostate cancer cells were seeded at optimal density in 384-well plates using an automated dispensing system (BioTek EL406). EZH2 inhibitors (GSK126 or EPZ-6438) were subjected to a 10-point series of threefold dilution (from 0.632 nM to 20 uM) in DMSO and then added into cells by robotic pin transfer in a JANUS workstation. Each drug at a certain dose in every specific cell line had four replicates. After 7 days of incubation, cellular ATP levels were measured using ATPlite Luminescence Assay (PerkinElmer). Data were normalized to the number of cells under DMSO conditions, and IC50 were determined with GraphPad Prism software.

### Standard ChIP and ChIP-Seq assays

Chromatin immunoprecipitation (ChIP) experiments were performed as previously described (Xu et al., 2012). Basically, cells were crosslinked with 1% formaldehyde and lysed in RIPA buffer with 0.3 M NaCl. ChIP DNA was purified using PCR Purification Kit (Qiagen) and then quantified by Quant-iTTM dsDNA HS Assay Kit (Invitrogen). Equal amounts of ChIP enriched DNA (5-10 ng) under each treatment condition (DMSO, GSK126 or EPZ-6438) were prepared for either targeted ChIP-qPCR or ChIP-Seq libraries. For protein detection in ChIP samples, SDS sample buffer was added to the reverse crosslinked input lysates, which were then subjected to Western blot analysis. ThruPLEX-FD Prep Kit (Rubicon Genomics) was used to construct the sequencing libraries according to the manufacturer’s protocol, and the final products were sequenced on the NextSeq 500. For targeted ChIP-qPCR, purified ChIP DNA was subjected to real-time quantitative PCR with specific primers as listed in Supplementary Table 2.

### ChIP-Rx of H3K27me3 in Prostate Cancer Cells

To quantitatively measure the changes of H3K27me3 abundance upon EZH2 inhibitor treatments, H3K27me3 ChIP with reference exogenous genome (ChIP-Rx) was performed as described (Orlando et al., 2014). Briefly, 5 ug of ready-to-ChIP human chromatin from abl or DU145 cells were mixed with 125 ng of Drosophila chromatin that has been sheared to proper sizes. 4 uL of H3K27me3 antibody together with 0.2 uL of Drosophila-specific H2Av antibody were added to the mixture. Each sample was then treated as one and subjected to standard processes of ChIP and sequencing. Short reads obtained from the sequencer were mapped to human genome (hg19) and drosophila genome (dm3) respectively, and peaks were called using MACS v2.0 (Zhang et al., 2008). Overall, 12,114 and 131,469 peaks of H3K27me3 were identified in abl and DU145 cells under DMSO treatment condition based on FDR<0.01. Normalization ratios between treatment and control groups were calculated based on the H2Av read counts mapped to drosophila genome between EPZ-6438 and DMSO or GSK126 and DMSO. The enrichment changes of human H3K27me3 signals were then normalized by the corresponding normalization ratios.

### Cell Transfections

A total of 50 pmol (for each well in 24-well plate) or 100 pmol (for each well in 6-well plate) of each siRNA was transfected into abl or DU145 cells using Lipofectamine RNAiMAX reagent (Invitrogen) according to the manufacturer’s instructions. Cell from 24-well plates were collected at indicated time points and counted after Trypan Blue staining for cell numbers, or harvested 48 hrs after transfection for RNA extraction, or lysed 72 hrs post-transfection and subjected to Western blot.

### RNA isolation and RT-qPCR

RNA was extracted and purified using the TRIzol Reagent combined with RNeasy Mini Kit (Qiagen) according to manufacturer’s protocols. 2 ug of total RNAs were then used for cDNA synthesis using High Capacity cDNA Reverse Transcription Kit (Applied Biosystems). Real-time quantitative RT-PCR was performed, and gene expression was calculated as described previously (Xu et al., 2012), using the formula 2-ΔΔCt relative to the level of GAPDH as the internal control. Sequences of RT-qPCR primers were listed in Supplemental Table 3.

### Cell Cycle Analysis with Flow Cytometry

Prostate cancer cells were pre-treated with nocodazole (5 ug/mL) for 24 hrs, and were released by being replenished with fresh medium. Cells were then incubated with GSK126 or EPZ-6438 at final concentrations of 5 uM for days as indicated. Cell cycle analyses were performed using previously published protocols. Generally, cells were collected, washed with ice-cold PBS, and fixed in 70% ethanol for at least 1 hr on ice. Cells were then pelleted, washed with PBS, and incubated in propidium iodide solution (Sigma, P4864) with RNase A (Sigma, R6513) for 30 min at 37°C. Flow cytometry analyses were done using an LSRII flow cytometer (Becton Dickinson).

### Comet Assay

Prostate cancer cells were treated under described conditions and the Alkaline Comet Assay was then performed following the manufacturer’s instructions (Trevigen, #4250-050-K). Briefly, treated or untreated cells were harvested and resuspended in ice cold PBS (Ca2+- and Mg2+-free) at a density of 1 x 10^5^ cells/ml, mixed with molten LMAgarose (1:10 ratio) and 50 μl of the mixture was immediately pipetted onto Comet Slide. After the agarose was solidified at 4°C in the dark for 15 minutes, the slides were immersed first in Lysis Solution for 1hr and then in Alkaline Unwinding Solution (200 mM NaOH, 1 mM EDTA, pH>13) for another 1 hr at 4°C in the dark. Alkaline electrophoresis was conducted at 4°C for 30 minutes at 21 volts. Cells were then fixed with 70% ethanol and stained with SYBR Gold. Comet images were taken by Nikon Ni-U fluorescence microscopy (Nikon) using a FITC filter. Comet tail moments were assessed using CometScore.v2.0 (TriTek Corp., Sumerduck, VA 22742, USA). Olive tail moment is defined as Tail DNA% x Tail Moment Length that is measured from the center of the head to the center of the tail. The quantification of Olive tail moments from each condition was calculated from a minimum of 100 cells for each data point.

### Test of EZH2 Inhibitors in Xenograft Mouse Model

Male nude mice (Taconic) were castrated after 3 days of accommodation. CWR22Rv1 xenografts were established in the flanks of mice by injecting ~2 million cells in 50% matrigel (BD Biosciences). When tumors reached approximately 200mm^3^, mice started to receive daily injection of EZH2 inhibitors (provided by Xcessbio Biosciences Inc) dissolved in 20% captisol (CYDEX, NC-04A-120106). Tumors were measured 3 times every week and harvested after 3 weeks. Frozen samples were subjected to RT-qPCR and immunoblotting to study the expression of selected genes. All animal protocols were approved by the Beth Israel Deaconess Institutional Animal Care and Use Committee, and the experiments were performed in accordance with institutional and national guidelines.

### Data Collection

EZH2 ChIP-Seq data and EZH2 siRNA microarray expression data were both retrieved from our previous study (GSE39461) (Xu et al., 2012). The Cancer Therapeutics Response Portal (CTRP) compound screen data was used in Figure 7 in order to correlate gene expression with cell sensitivity to EZH2 inhibitors (Rees et al., 2016). Two independent prostate cancer cohorts were retrieved and analyzed in Figure 5F and G (Taylor et al., 2010) or Figure S7D and E (Grasso et al., 2012). In Figure S7F, the LNCaP RNA-Seq data sets were downloaded from the published data sets (Chen et al., 2015).

### Analysis of EZH2 Inhibitor-Mediated Gene Expression by RNA-Seq

Both abl and DU145 cells were treated with EZH2 inhibitors (GSK126 or EPZ-6438) at final concentrations of 5 uM for 60-72 hrs before RNAs were extracted. RNA-seq library was prepared using Illumina True-seq RNA sample preparation kit and sequenced to 50bp using Illumina Hi-seq platform. RNA-seq data was mapped to human genome (hg19) using TopHat version 2.0.6 (Trapnell et al., 2009). DESeq2 was applied to calculate the logarithmic fold change (LFC) and p-value in order to call any significantly changed genes between treatment and control groups. Differentially expressed genes were first filtered using LFC >0.5 or <-0.5, and then top 200 genes were selected through ranking by their p-values. The authentic target genes of EZH2 in abl cells were defined as those showing similar expression changes upon either EZH2 silencing or inhibitor treatment.

### Focused CRISPR screen library design

To design a smaller scale CRISPR/Cas9 knockout screen library focusing on cancer-related genes, we selected 6000 genes based on the reported literatures (Forbes et al., 2017; Garcia et al., 2017; Huether et al., 2014; Lawrence et al., 2014; Vogelstein et al., 2013). For each gene, we designed ten single-guide RNAs (sgRNAs) with 19bp against its coding region with optimized cutting efficiency and minimized off-target potentials. Cutting efficiency wise, we used sequence features of the spacers to calculate the efficiency score for each sgRNA using predictive model (Xu et al., 2015). Off-target wise, we used BOWTIE to map all candidate sgRNAs to hg38 reference genome, and chose those with least potential off-targets (Langmead et al., 2009). We selected the 10 best sgRNAs for each gene based on the criteria above. The library also contains positive and two types of negative controls (nontargeting controls and non-essential regions-targeting sgRNAs).

1. Positive controls: we included 1466 sgRNAs targeting 147 positive control genes, which are significantly negatively selected in multiple screen conditions.
2. Non-targeting negative controls: 795 sgRNAs with sequences not found in genome.
3. Non-essential regions-targeting negative controls: 1891 sgRNAs targeting AAVS1, ROSA26, and CCR5, which have been reported as safe-harbor regions where knock-in leads to few detectable phenotypic and genotypic changes (Sadelain et al., 2011).

### Plasmid construction and lentivirus production

The sgRNA library was synthesized at CustomArray© and then amplified by PCR. The PCR products were subsequently ligated into lentiCRISPR V2 plasmid, followed by transformation into competent cells according to an online protocol (GeCKO library Amplification Protocol from Addgene). Afterwards, we isolated the plasmid and constructed a sequencing library for Miseq to ensure library diversity. To make lentivirus, T-225 flasks of 293FT cells were cultured at 40%~50% confluence the day before transfection. Transfection was performed using X-tremeGENE HP DNA Transfection Reagent (Roche). For each flask, 20 ug of lentivectors, 5 ug of pMD2.G, and 15 ug of psPAX2 (Addgene) were added into 3 ml OptiMEM (Life Technologies). 100 ul of X-tremeGENE HP DNA Transfection Reagent was diluted in 3 ml OptiMEM and, after 10 min, it was added to the plasmid mixture. The complete mixture was incubated for 20 min before being added to cells. After 6 hr, the media was changed to 30 ml DMEM + 10% FBS. The media was removed 60 hrs later and centrifuged at 3,000 rpm at 4 °C for 10 min to pellet cell debris. The supernatant was filtered through a 0.45 um membrane with low protein binding. The virus was ultracentrifuged at 24,000 rpm for 2 hr at 4 °C and then resuspended overnight at 4°C in DMEM + 10% FBS. Aliquots were stored at −80°C.

### CRISPR screens

LNCaP or abl were kept in their respective medium that is routinely used to maintain normal proliferation. Cells of interest were infected at a low MOI (0.3~0.5) to ensure that most cells receive only 1 viral construct with high probability. Briefly, 3×10^6^ cells per well were plated into a 12 well plate in the appropriate standard media supplemented with 8ug/ml polybrene. Each well received a different titrated virus amount, usually between 5 and 50 ul, along with a non-transduction control. The 12-well plate was centrifuged at 2,000 rpm for 2 hr at 37°C. After the spin, media was aspirated and fresh media without polybrene was added. Cells were incubated overnight and then enzymatically detached using trypsin. Cells were counted and each well was split into duplicate wells. Each replicate of LNCaP or abl cells received 4ug/mL puromycin. After 3 days or as soon as no surviving cells remained in the non-transduction control under puromycin selection, cells were counted. Percent transduction was calculated as cell numbers from the replicate with puromycin divided by cell counts from the replicate without puromycin and then multiplied by 100. The virus volume yielding a MOI closest to 0.4 was chosen for large-scale screening.

For focused CRISPR-Cas9 knockout screen, large-scale spin-infection of 2×10^8^ cells was carried out using four of 12-well plates with 4×10^6^ cells per well. Wells were pooled together into larger flasks on the same day after spin-infection. After three days of puromycin selection, the surviving abl cells were divided into three groups: one for day 0 control, and the other two cultured in the presence of DMSO or GSK126 for four weeks. Two rounds of PCR were performed after gDNA had been extracted, and 300ug DNA per sample was used for library construction. Each library was sequenced at 3~30 million reads to achieve ~300X average coverage. The day 0 sample library served as the control to identify genes or pathways that were positively or negatively selected.

### CRISPR screen normalization and analysis

We analyzed CRISPR knockout screen data following MAGeCK protocol, which assigns each gene with a beta score (β), analogy of log fold change (Li et al., 2015). Positive and negative beta scores indicate positive and negative selection, respectively. Considering that cells in screens may be harvested at different time points, normalization of screen duration was carried out to ensure comparable CRISPR screens. Equation of cell growth is equivalent to that of beta score defined in MAGeCK:

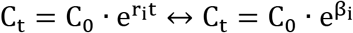

Where r_i_ is growth constant of cells with gene g_i_ knockout, and t is the duration of screen. The equivalence of these two equations shows that beta scores are linearly dependent on screen duration, suggesting the necessity of equalizing the screen durations for fair screen comparisons. Considering that the pan-essential genes are negatively selected similarly in different conditions, the absolute median beta score of the pan-essential genes is proportional to the screen duration. Assuming the r of essential genes remains constant in various screen conditions, then:

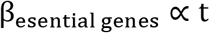

Therefore, we can rescale the beta scores using the absolute median beta scores of known-essential genes (Hart et al., 2014):

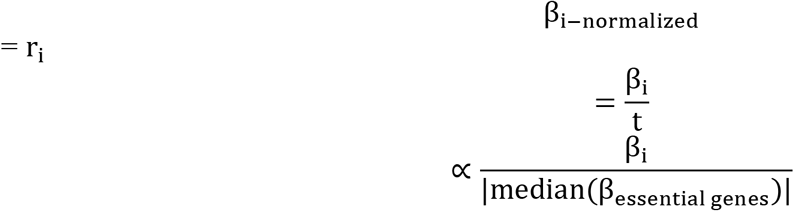

## Declaration of Interests

N.S. receives research grant funding from AstraZeneca.

J.B. is an executive and shareholder of Novartis AG, and has been a founder and shareholder of SHAPE (acquired by Medivir), Acetylon (acquired by Celgene) Tensha (acquired by Roche), Syros, Regency and C4 Therapeutics.

P.K. serves on the SAB of BIND Biosciences, BN Immunotherapeutics, GE Healthcare, Janssen, New England Research Institutes, OncoCellMDX, Progenity, Sanofi and Thermo Fisher. He shares investment interests in Context Therapeutics, DRGT, Placon, Seer Biosciences and Tarveda Therapeutics. He also serves on the DSMB of Genetch and Merck.

X.S.L. is a cofounder and board member of GV20 Oncotherapy, SAB of 3DMed Care, consultant for Genentech, and stockholder of BMY, TMO, WBA, ABT, ABBV and JNJ.

M.B. serves on the SAB of Kronos Bio and is a consultant to Aleta BioTherapeutics and H3 Biomedicine.

All the other authors declare no potential conflicts of interest.

## Acknowledgement

The authors sincerely thank Dr. Matthew Freedman for sharing their normal prostate epithelial cell lines. We are also very grateful to Yin Liu for technical support. Finally we want to thank Anna Groner, Housheng Hansen He and Prakash K. Rao for their insightful advice and discussion. This work was supported by grants (HG008927 to X.S.L., CA090381 and CA163227 to M.B. and R00 CA178199 to K.X.) from National Institutes of Health, funding supports (PC140817P1 to M.B. and W81XWH-16-1-0409 to N.S.) from the Department of Defense, AACR-AZ START Grant (19-40-12 to N.S.), Prostate Cancer Foundation Young Investigator award (to K.X.), CPRIT award (RR140072 to K.X.), and Voelcker Fund Young Investigator award (to K.X.).

## Accession Number

All the genome-wide datasets generated in this study, including RNA-Seq of EZH2 inhibitors and ChIP-Rx of H3K27me3, were deposited at the Gene Expression Omnibus database (http://www.ncbi.nlm.nih.gov/geo/) with an accession number GSE80240.

## Author Contributions

Y.L., N.S., M.Y., J.L., T.F., M.D., J.Z. and K.X. performed all the biochemical, biological and molecular biology assays in human prostate cells, ChIP-Seq, RNA-Seq. C.C., A.F., H.X., W.L., S.M. and X.S.L did all the genomic data analyses, clinical data statistical significance calculation and synergy score calculation. T.X. carried out the CRISPR-Cas9 knockout screens. C.C., S.G. and S.B. established the xenograft models and tested the efficacy of EZH2 inhibitors in vivo. P.X. and Z.L. helped with cell culture and Comet assays. M.M., R.P., S.S., J.B., and K.P. assisted with drug sensitivity screens in vitro. This study was conceptually monitored by X.S.L, M.B. and K.X. with the help from P.K., H.L. and S.B. All authors helped design the study and write the manuscript.

## Notes

### Competing Interest Statement

The authors have declared no competing interest.

