## Supplemental Information for "A non-canonical EZH2 function sensitizes solid tumors to genotoxic stress"

CONTENTS:

Supplemental Figures

Figure S1, related to Figure 1

Figure S2, related to Figure 1

Figure S3, related to Figure 2

Figure S4, related to Figure 2

Figure S5, related to Figure 3

Figure S6, related to Figure 4

Figure S7, related to Figure 5

Figure S8, related to Figure 6

Figure S9, related to Figure 7

Supplemental Tables

Table S1. Prostate cell lines and their culture conditions

Table S2. Primers for quantitative real-time RT-PCR

Table S3. Primers for targeted ChIP-qPCR

Supplemental References

Supplemental Figures

Fig. S1, related to Fig. 1

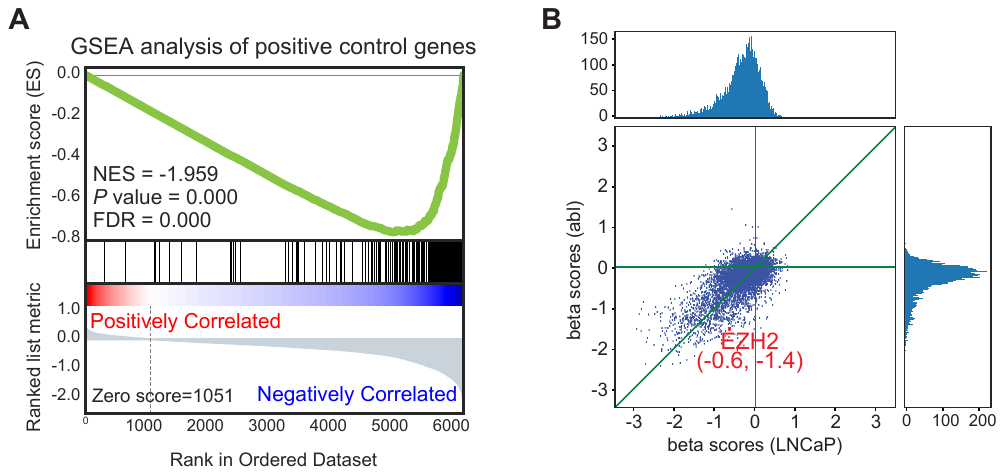

**Fig. S1. Quality control of CRISPR-Cas9 knockout screening in LNCaP and abl cells.**

**(A).** Gene set enrichment analysis (GSEA) of positive sgRNAs that are known to regulate cell proliferation showed that cell clones containing these sgRNAs were indeed negatively selected.

**(B).** Distributions of beta scores from the CRISPR-Cas9 knockout screening in LNCaP (x-axis) and abl (y-axis) cells were plotted. Position of EZH2 was indicated by red dots. Numbers in the parentheses, beta scores in LNCaP (the first values) and abl (the second values).

Fig. S2, related to Fig. 1

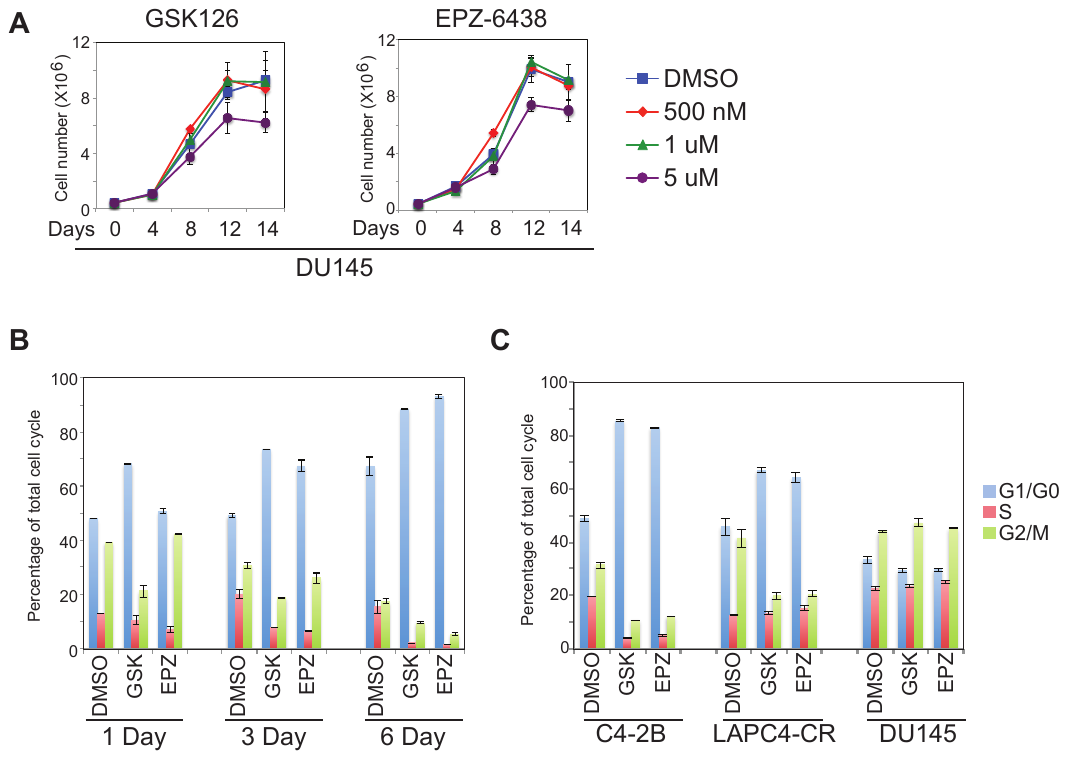

**Fig. S2. Effects of EZH2 inhibitors on the proliferation and cell cycle of prostate cancer cells.**

**(A).** Effects of EZH2 inhibitors (GSK126 and EPZ-6438) on the growth of DU145 cells over time with indicated concentrations of the compounds.

**(B).** Cell cycle analysis of abl cells with the treatment of vehicle (DMSO), 5 uM GSK126 (GSK) or 5 uM EPZ-6438 (EPZ) over indicated days.

**(C).** Prostate cancer cells were treated with vehicle (DMSO), 5 uM GSK126 (GSK) or 5 uM EPZ-6438 (EPZ) for 3 days, and then subjected to propidium iodide staining followed by flow cytometry analysis.

Fig. S3, related to Fig. 2

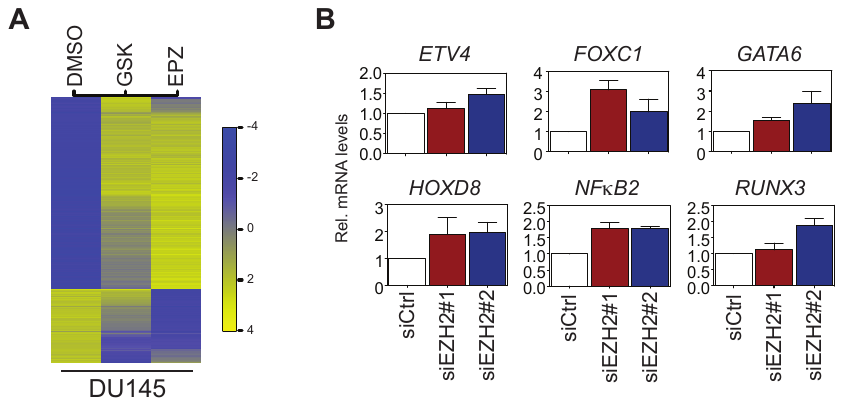

**Fig. S3. EZH2-regulated gene expression profiles in irresponsive prostate cancer cells.**

**(A).** Heat map of differential genes in DU145 cells, which were treated with vehicle (DMSO), 5 uM GSK126 (GSK) or 5 uM EPZ-6438 (EPZ) for 72 hrs.

**(B).** DU145 cells were transfected with control siRNA (siCtrl), or two independent siRNAs specific for EZH2 (siEZH2#1 and #2). Total RNA was extracted 48 hrs after transfection, and real-time RT-qPCR was performed to detect the changes in mRNA levels of indicated genes.

Fig. S4, related to Fig. 2

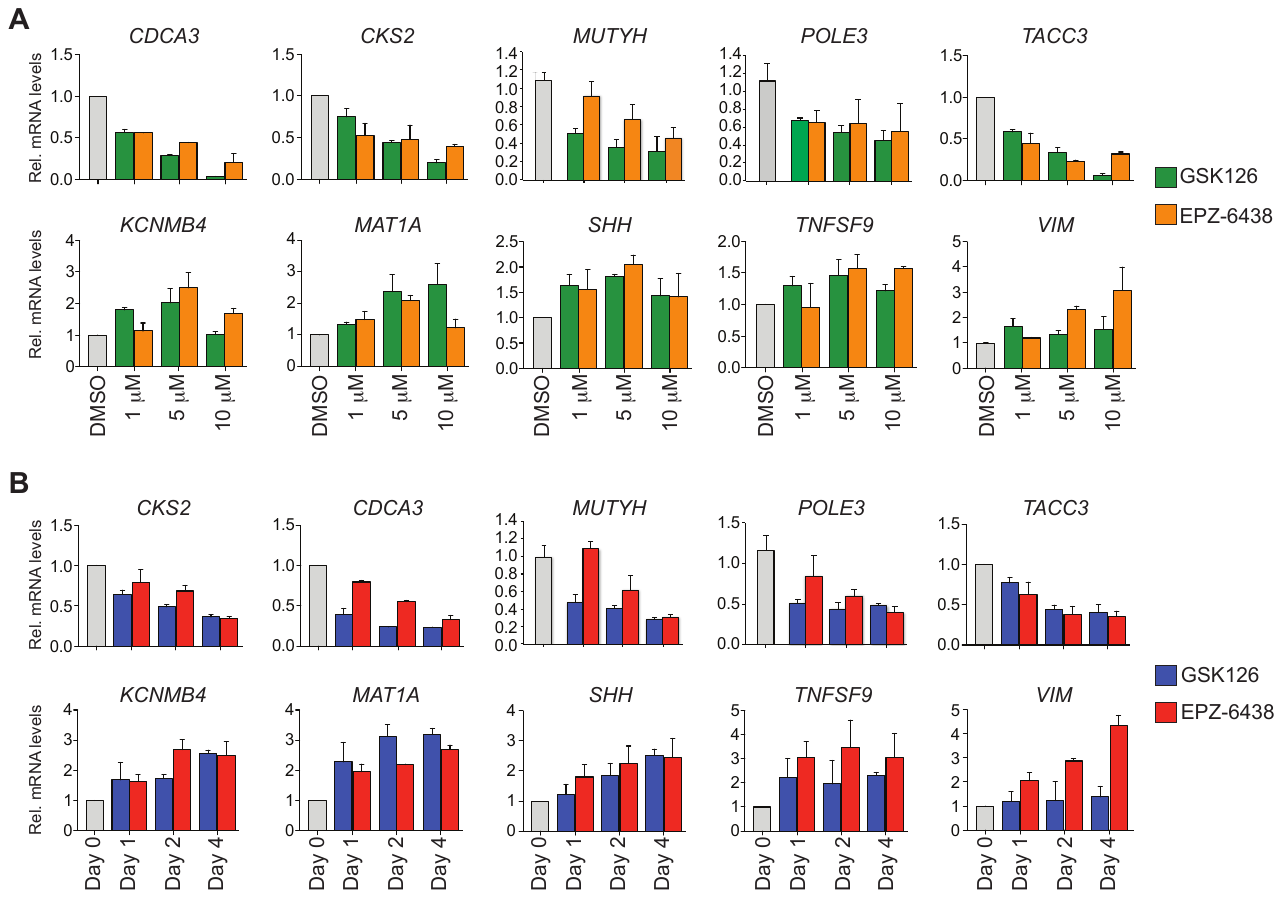

**Fig. S4. Confirmation of EZH2 inhibitor-induced gene expression patterns in sensitive prostate cancer cells.**

**(A-B).** Abl cells were treated with GSK126 or EPZ-6438 at different concentrations for 72 hrs (A) or at 5 uM for indicated days (B). Real-time RT-qPCR was carried out to examine the expression of selected genes.

Fig. S5, related to Fig. 3

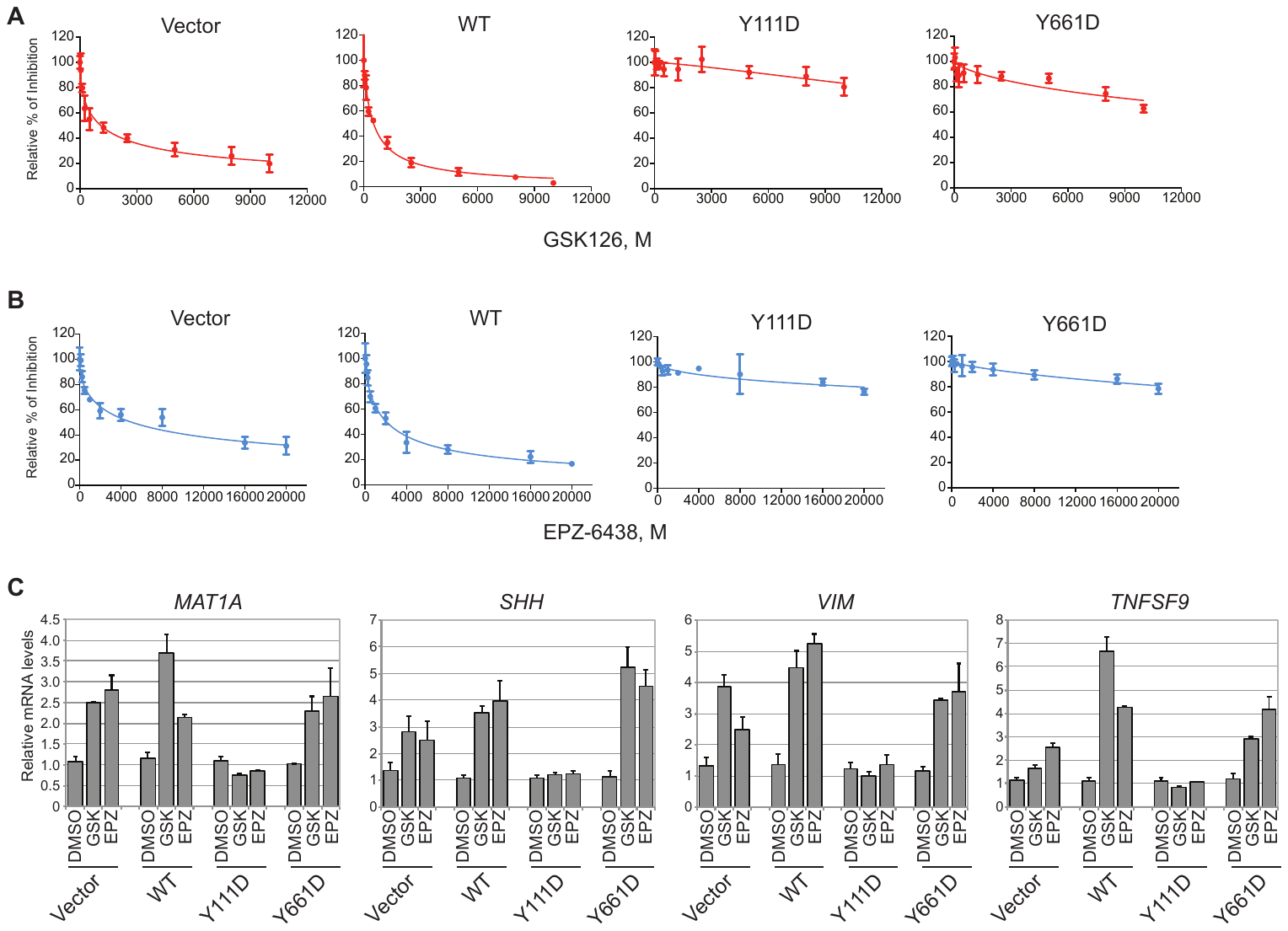

**Fig. S5. Resistance of Y111D- and Y661D-expressing abl cells to EZH2 inhibitors.**

**(A-B).** Abl cells expressing the control (Vector), the wild-type EZH2 (WT), or mutants EZH2 (Y111D and Y661D) were incubated with indicated concentrations of GSK126 (A) or EPZ-6438 (B) for 6 days. Cell numbers were counted at the end point after trypan blue staining, and normalized to that under vehicle condition, which was considered as 100%.

**(C).** Abl cells expressing the control (Vector), the wild-type EZH2 (WT), or various EZH2 mutants (Y111D and Y661D) were treated with vehicle (DMSO) or 5 μM EZH2 inhibitors (GSK, GSK126; EPZ, EPZ-6438) for 72 hrs. Afterwards, total RNA was extracted and RT-qPCR was followed to detect the transcript levels of indicated genes. The levels of *GAPDH* were served as the internal control.

Fig. S6, related to Fig. 4

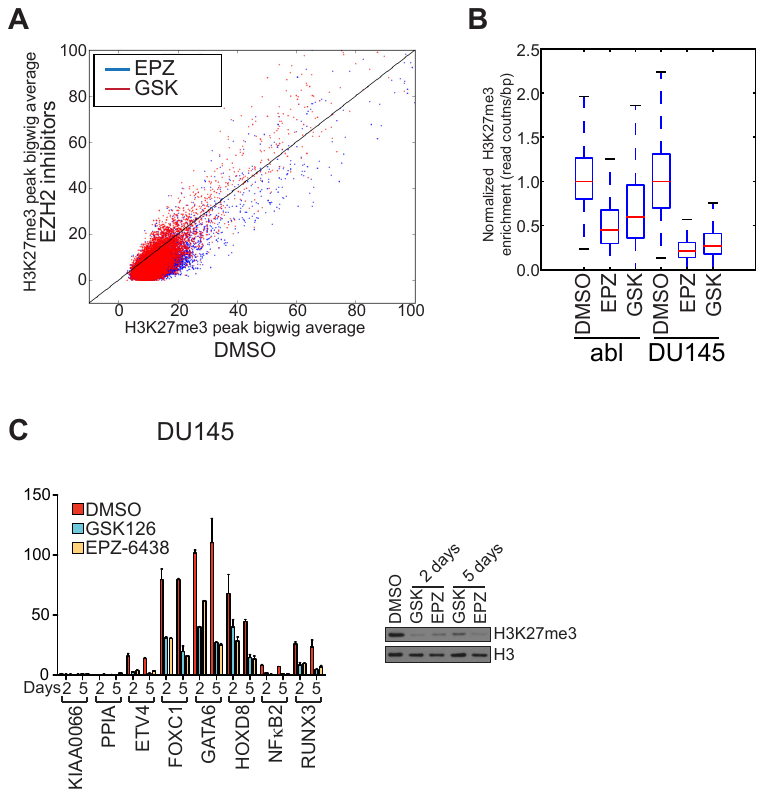

**Fig. S6. Canonical method of normalizing genome-wide H3K27me3 signals in prostate cancer cells upon the treatment of EZH2 inhibitors.**

**(A).** Intensity of each H3K27me3 peak was plotted and compared between control condition (DMSO, x-axis) and treatment condition (EZH2 inhibitors, y-axis) in abl cells, using reads per million methods.

**(B).** H3K27me3 peak enrichment under the conditions of vehicle (DMSO), 5 uM GSK126 (GSK) or 5 uM EPZ-6438 (EPZ) was normalized using canonical method and then compared between abl and DU145 cells.

**(C).** Direct ChIP-qPCR of H3K27me3 was performed and detected at selected chromatin regions in DU145 cells. Cells were treated with vehicle (DMSO) or EZH2 inhibitors for indicated number of days. KIAA0066 and PPIA, negative controls; right panel, H3K27me3 protein levels by immunoblotting in the corresponding ChIP samples. GSK, GSK126; EPZ, EPZ-6438.

Fig. S7, related to Fig. 5

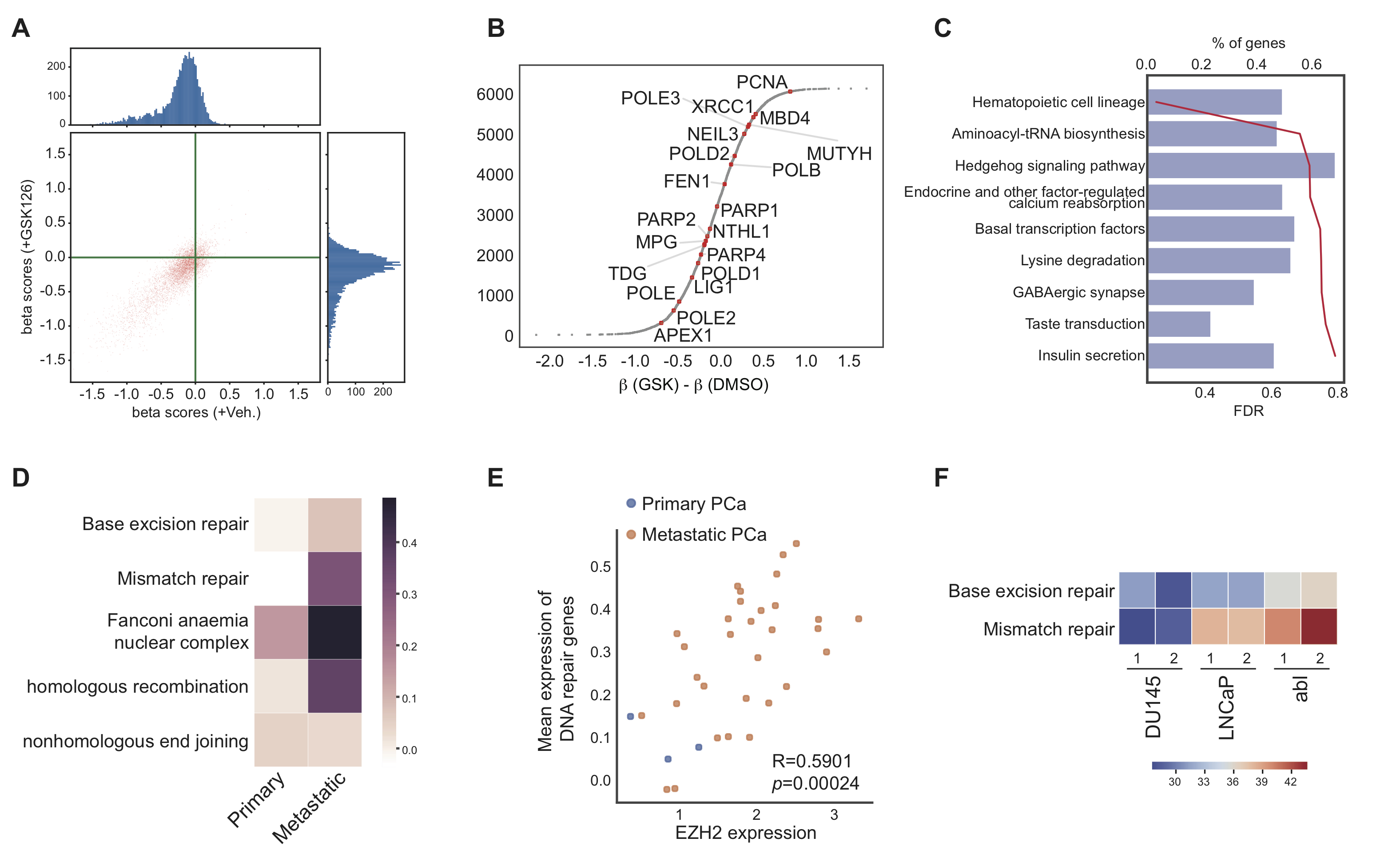

**Fig. S7. Identification of signature genes that are essential for the effects of EZH2 inhibitors in prostate cancer cells.**

**(A).** Distributions of beta scores from the CRISPR-Cas9 knockout screening in LNCaP cells were plotted for both control condition (+Veh., x-axis) and treatment condition (+GSK126, y-axis).

**(B).** Distribution of delta beta scores in LNCaP cells, defined as beta scores under treatment (+GSK126) condition minus beta scores under control (+Veh.) condition. Positive values of beta scores indicate that the genes show weaker essentiality in the presence of GSK126 than in the presence of DMSO. Representative DNA repair genes were marked by red dots.

**(C).** Functional annotations that were enriched in genes with positive delta beta scores from CRISPR-Cas9 knockout screening in LNCaP cells were shown. Blue bars, percentage of genes in each specific functional category; red line, values of the false discovery rate (FDR) for the particular gene ontology term.

**(D).** Heat map showing expression of DNA damage repair genes in another independent prostate cancer cohort containing primary or metastatic tumors (*1*). Genes were categorized into different DNA repair pathways according to their functions.

**(E).** Expression correlation between EZH2 and DNA damage repair genes was plotted based on another independent prostate cancer cohort (*1*). Each dot represents a clinical case, and all the primary prostate cancer (PCa) were colored in blue, while the metastatic ones were colored in yellow.

**(F).** The transcript levels (TPM) of EZH2-activated genes that are functionally involved in either base excision repair or mismatch repair pathway were compared among LNCaP, abl and DU145. Numbers, duplicates of data was utilized in the analysis.

Fig. S8, related to Fig. 6

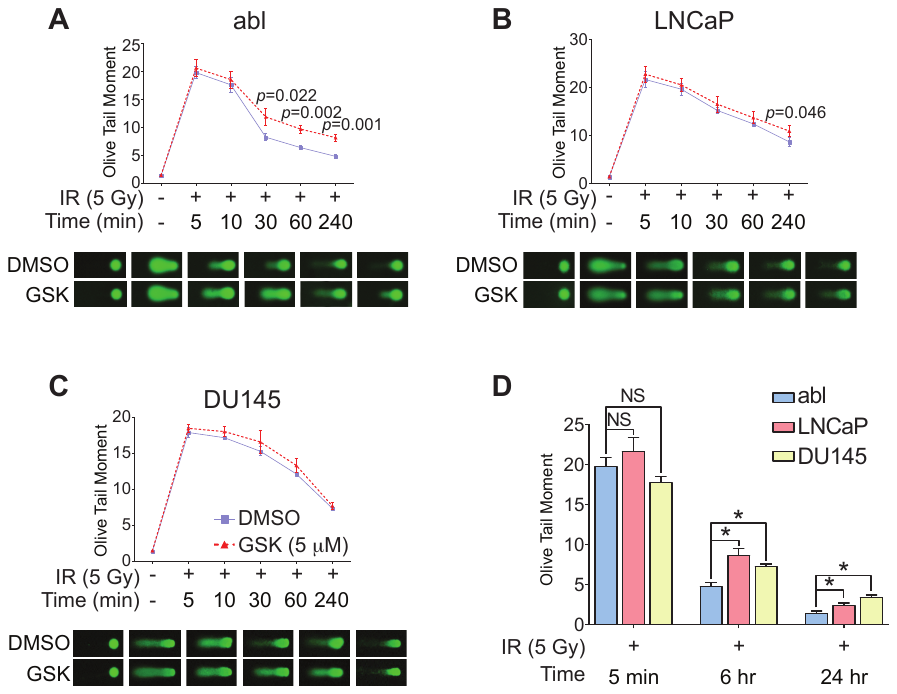

**Fig. S8. Responses of prostate cancer cells to DNA damage agents in the presence of EZH2 inhibitors.**

**(A-C).** Alkaline comet assays were carried out in abl (A), LNCaP (B) or DU145 (C). Prostate cancer cells were pretreated with vehicle (DMSO) or 5 µM GSK126 (GSK) for 7 days, and then exposed to 5 gray (Gy) ionizing radiation (IR) followed by recovery at indicated time points. Top panels, quantification of the results using “Olive Tail Moment” parameter; bottom panels, representative images of each time point.

**(D).** Abl, LNCaP and DU145 were treated with 5 gray (Gy) ionizing radiation (IR) followed by recovery at indicated time points. The basal efficiency of DNA repair in each cell line was plotted and compared using “Olive Tail Moment” parameter; *, *p*<0.01.

Fig. S9, related to Fig. 7

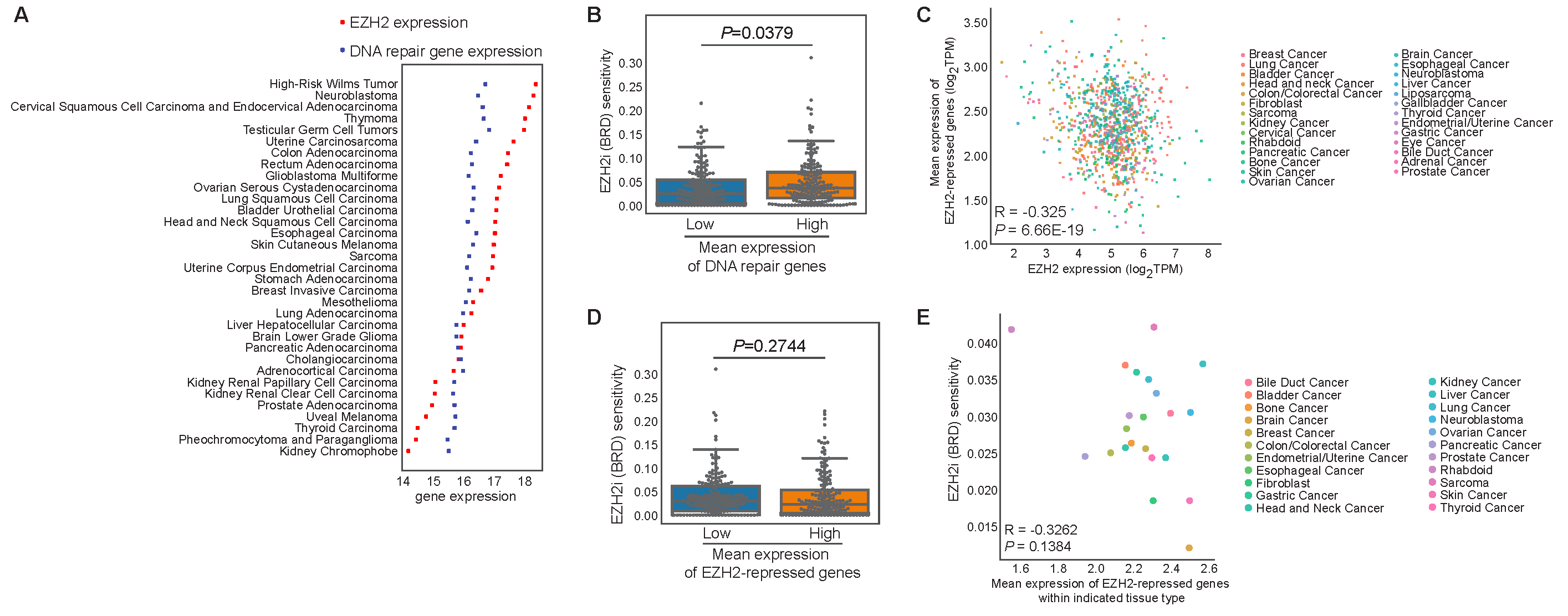

**Fig. S9. Correlation between expression of DNA damage repair genes and responses of various types of cancer cells to EZH2 inhibitors.**

**(A).** Expression levels of EZH2 (red dots) and DNA repair genes (blue dots) in each specified cancer type in TCGA data (*2*).

**(B and D).** Box plots showing the sensitivity of CTRP cell lines that do not contain EZH2 mutations to EZH2 inhibitor (BRD, BRD-K62801835-001-01-0) (*3*). Cells were grouped based on mean expression of DNA repair genes (B) or EZH2-repressed genes (D), and then sensitivities to BRD in the top 25% of cells with the highest expression were compared with those in the bottom 25% with the lowest expression.

**(C).** Expression correlation between EZH2-repressed genes and EZH2 in cancer cells from CCLE gene expression data (*4*). Each dot represents one cell line and those within the same type were colored with the same color.

**(E).** Dot plots demonstrating the association between expression of EZH2-repressed genes and sensitivity to EZH2 inhibitor within specified tissue types. Cellular sensitivity to EZH2 inhibitor (BRD, BRD-K62801835-001-01-0) was derived from CTRP compound screen data (*3*).

Supplemental Tables

**Table S1. Prostate cell lines and their culture conditions.**

| Cell Names | Culture Medium | Supplements | Culture Condition |
| --- | --- | --- | --- |
| Prostate epithelial cell lines | | | |
| LHSAR | PrEBM basal medium | PrEGM SingleQuot Kit Suppl. & Growth Factors | 37°C, 5% CO2 |
| RWPE-1 | Keratinoyte serum free medium | 0.05 mg/mL bovine pituitary extract (BPE), 5 ng/mL human recombinant epidermal growth factors (EGF) | 37°C, 5% CO2 |
| AR-null prostate cancer cell lines | | | |
| DU145 | phenol-red-free RPMI1640 | 10% charcoal-stripped FBS, 1% Penicillin-Streptomycin | 37°C, 5% CO2 |
| PC3 | regular DMEM | 10% FBS, 1% Penicillin-Streptomycin | 37°C, 5% CO2 |
| AR-positive, androgen-dependent prostate cancer cell lines | | | |
| LAPC4 | regular RPMI1640 | 10% FBS, 1% Penicillin-Streptomycin | 37°C, 5% CO2 |
| LNCaP | regular RPMI1640 | 10% FBS, 1% Penicillin-Streptomycin | 37°C, 5% CO2 |
| VCaP | regular DMEM | 15% FBS, 1% Penicillin-Streptomycin, 1% Non-essential amino acids | 37°C, 5% CO2 |
| AR-positive, androgen-independent prostate cancer cell lines | | | |
| C4-2B | regular RPMI1640 | 10% FBS, 1% Penicillin-Streptomycin | 37°C, 5% CO2 |
| CWR22Rv1 | regular DMEM | 10% FBS, 1% Penicillin-Streptomycin | 37°C, 5% CO2 |
| LAPC4-CR | phenol-red-free RPMI1640 | 10% charcoal-stripped FBS, 1% Penicillin-Streptomycin | 37°C, 5% CO2 |
| LNCaP-abl | phenol-red-free RPMI1640 | 10% charcoal-stripped heat-inactivated FBS, 1% Penicillin-Streptomycin | 37°C, 5% CO2 |
| LNCaP-AI | phenol-red-free RPMI1640 | 10% charcoal-stripped FBS, 1% Penicillin-Streptomycin | 37°C, 5% CO2 |

**Table S2. Primers for targeted ChIP-qPCR.**

| Names | Sequences |
| --- | --- |
| KIAA0066 F | CTAGGAGGGTGGAGGTAGGG (*5*) |
| KIAA0066 R | GCCCCAAACAGGAGTAATGA (*5*) |
| PPIA F | GCCAGGCTCCTGTTTTAATG |
| PPIA R | GCAGTCTCCGGTTTTGAGAG |
| CCND2 F | TCCAACCGAAACTCCAAAAC (*6*) |
| CCND2 R | CTTTTCACCCTTCACGGAAA (*6*) |
| DAB2IP F | CCTGCTCTGAGTCTGCACTG (*6*) |
| DAB2IP R | TCGAATCTCTCCCATGGTTC (*6*) |
| p16 F | AGGGGAAGGAGAGAGCAGTC (*7*) |
| p16 R | GGGTGTTTGGTGTCATAGGG (*7*) |
| SHH F | TCCTTCCATTTCCACTCCTG |
| SHH R | TCTTGCTACAATGGCCTTCC |
| TNFSF9 F | GCGATTTCTTGGCGTTACTT |
| TNFSF9 R | TCGGGGAGGTTAGAGTGCT |
| VIM F | CAATCTCAGGCGCTCTTTGT |
| VIM R | GAGCGGGAAGAGGAAAGAGTA |
| ETV4 F | TCTCCAGCCTATGCACTCCT |
| ETV4 R | CTTCCATTTGCACAAGCAGA |
| FOXC1 F | CCCTCTCTTGCCTTCTTCCT |
| FOXC1 R | CGTCAGGTTTTGGGAACACT |
| GATA6 F | CCCTAACTGGGAAAACACGA |
| GATA6 R | CGCCCAGGTAAATCCAAGTA |
| HOXD8 F | AATAGTTCGGGTGCGTTTTG |
| HOXD8 R | TCACTGGCCCAATCTTTTTC |
| NFkB2 F | GGGGTGGGGAAGTAATAGGA |
| NFkB2 R | CCTTAGCAGGTGCCATGAGT |
| RUNX3 F | CATGGACCGTAGTCTTTTCT |
| RUNX3 R | CACTGCCAAGAACGCACTTA |

**Table S3. Primers for quantitative real-time RT-qPCR.**

| Gene Names | Sequences |
| --- | --- |
| TACC3 mRNA F | GCACAGGATTCTAAGTCCTAGCA |
| TACC3 mRNA R | CCAGACCGGGTGTGAGTTTT |
| KIAA0101 mRNA F | ATGGTGCGGACTAAAGCAGAC (*6*) |
| KIAA0101 mRNA R | CCTCGATGAAACTGATGTCGAAT (*6*) |
| BIRC5 mRNA F | AGGACCACCGCATCTCTACAT |
| BIRC5 mRNA R | AAGTCTGGCTCGTTCTCAGTG |
| CDCA3 mRNA F | CTGGAGGGTCTTAAACATGCC |
| CDCA3 mRNA R | CACTGCTGGTCTTCATAGGTG |
| SHH mRNA F | CCAAGGCACATATCCACTGCT |
| SHH mRNA R | GTCTCGATCACGTAGAAGACCT |
| VIM mRNA F | AGTCCACTGAGTACCGGAGAC |
| VIM mRNA R | CATTTCACGCATCTGGCGTTC |
| MAT1A mRNA F | ATCAGGGTTTGATGTTCGGCT |
| MAT1A mRNA R | GCGTTGAGCTTGTGAGCAA |
| TNFSF9 mRNA F | GGCTGGAGTCTACTATGTCTTCT |
| TNFSF9 mRNA R | ACCTCGGTGAAGGGAGTCC |
| CKS2 mRNA F | TTCGACGAACACTACGAGTACC (*6*) |
| CKS2 mRNA R | GGACACCAAGTCTCCTCCAC (*6*) |
| APEX1 mRNA F | GTTTCTTACGGCATAGGCGAT |
| APEX1 mRNA R | CACAAACGAGTCAAATTCAGCC |
| POLE mRNA F | TTCCTCAGTTTCGGCACTCAA |
| POLE mRNA R | CTCAAAACCAAACCGCAAATCC |
| POLE3 mRNA F | GCTGTACGCCACATCCTGT |
| POLE3 mRNA R | TTCAATGGGGTAACGAACCGC |
| MUTYH mRNA F | TGCCACGTACAGCAGAGAC |
| MUTYH mRNA R | CAAAGGCGATAGAGGCAATGG |
| FOXC1 mRNA F | TGTTCGAGTCACAGAGGATCG |
| FOXC1 mRNA R | ACAGTCGTAGACGAAAGCTCC |
| ETV4 mRNA F | GCAACGGAATTTCCTGAGATCC |
| ETV4 mRNA R | ACGGAGCTATGTTCCCCGA |
| GATA6 mRNA F | GTGCCAACTGTCACACCACA |
| GATA6 mRNA R | GAGTCCACAAGCATTGCACAC |
| HOXD8 mRNA F | GGAAGACAAACCTACAGTCGC |
| HOXD8 mRNA R | TCCTGGTCAGATAGGGGTTAAAA |
| RUNX3 mRNA F | AGCACCACAAGCCACTTCAG |
| RUNX3 mRNA R | GGGAAGGAGCGGTCAAACTG |
| NFB2 mRNA F | AGAGGCTTCCGATTTCGATATGG |
| NFB2 mRNA R | GGATAGGTCTTTCGGCCCTTC |
| KCNMB4 mRNA F | AGTGCTCCTATATCCCTCCCT |
| KCNMB4 mRNA R | GCTGGGAACCAATCTCATCTTT |
| GAPDH mRNA F | CGAGATCCCTCCAAAATCAA(*6*) |
| GAPDH mRNA R | TTCACACCCATGACGAACAT(*6*) |
